# Global Hypomethylation in Cell-free DNA Enables Non-invasive Colorectal Cancer Screening: Results from a Retrospective Validation Study

**DOI:** 10.64898/2025.12.14.694262

**Authors:** Shahdeep Kaur, Shefali Lathwal, Smita Agrawal, Tina Mahmoudi, Shalini Gupta

## Abstract

Global DNA hypomethylation is a defining hallmark of colorectal cancer (CRC) but is poorly captured by existing cell-free DNA (cfDNA) technologies, which typically interrogate only a fraction of CpG sites and are biased toward CpG islands. We developed Asima Rev, an electrical-impedance cfDNA assay that can differentiate healthy individuals from those with cancer by measuring cfDNA aggregation patterns associated with methylation state, enabling functional detection of genome-wide hypomethylation. In a cohort of 46 treatment-naive CRC patients and 33 controls, Asima Rev achieved 96% sensitivity and 94% specificity, with longitudinal monitoring in six patients fully concordant with clinical outcomes. Interrogation of 11 public methylation array datasets showed that whole array analyses underestimate global changes. However, restricting analyses to OpenSea regions, where cfDNA is enriched, revealed patterns consistent with Asima Rev and a significant 5.8% global hypomethylation in CRC tissue compared to adjacent normal tissue. Hypomethylation was not observed in immune cell genomic DNA (gDNA), a major contributor to cfDNA, supporting a predominantly tumor-derived contribution to the observed cfDNA signal. Together, these results demonstrate that Asima Rev captures a cfDNA signal consistent with true global methylation loss and outperforms locus-specific assays by measuring structural consequences of pan-genomic epigenetic alterations.

## Introduction

Colorectal cancer (CRC) is the third most common cancer globally and the second leading cause of cancer-related mortality, accounting for nearly 1 million deaths annually^1^. Early detection and timely intervention significantly improve survival, with 5-year rates exceeding 70% for locoregional disease^2^. Yet, population-level screening adherence remains suboptimal, particularly in low-resource or decentralized settings. Despite extensive public health campaigns, only about 60–65% of eligible individuals in the United States undergo routine CRC screening^3^, with even lower adherence in underserved populations. This gap underscores the need for non-invasive, scalable diagnostic tools that are clinically robust, operationally simple and economically viable.

Epigenetic alterations, particularly DNA methylation, are a hallmark of many cancers, including CRC^4^. Global hypomethylation was first described in CRC tissues more than four decades ago using genome-wide biochemical assays such as enzymatic digestion followed by liquid chromatography^5^ or methyl-cytosine-sensitive restriction endonucleases followed by gel electrophoresis^6^. While crude by modern standards, these methods truly captured whole-genome methylation loss. Newer technologies such as methylation arrays, sequencing, and polymerase-chain reaction (PCR)-based panels have confirmed that CRC tissues exhibit widespread global hypomethylation along with localized hypermethylation at promoter CpG islands^4,7^, but have primarily focused on site-specific changes^8^. Distinct epigenetic patterns also define CRC subtypes, including the CpG Island Methylator Phenotype (CIMP) characterized by vast hypermethylation of promoter CpG island sites^9^, microsatellite instability (MSI) driven by methylation-mediated mismatch repair silencing^10^, and consensus molecular subtypes (CMS1-4)^11^, each characterized by specific epigenetic and genomic alterations. This increased resolution has shifted clinical applications toward promoter-targeted methylation assays^4^.

Most modern methylation technologies interrogate only a small fraction of the genome. Methylation arrays typically cover 450k–850k CpGs (∼1.6–3.0% of the ∼28 million CpG sites in the human methylome)^12^ while sequencing-based assays often rely on restricted panels due to the high cost of whole-genome bisulfite sequencing. These single-nucleotide or short-region measurements - akin to examining individual “beads” on a DNA necklace - enable detailed molecular profiling but require complex workflows, specialized personnel, and costly instrumentation. As a result, broad population deployment remains challenging.

Cell-free DNA (cfDNA) has emerged as a minimally invasive analyte for CRC screening, prognosis, and disease monitoring. Yet, despite global hypomethylation being one of the earliest and most fundamental epigenetic hallmarks of CRC tissues, it remains underexplored in cfDNA. Technical barriers including high DNA input requirements for arrays (∼200-250 ng/test)^13,14^, sensitivity limitations, and the predominance of clinical interest in targeted gene panels have restricted cfDNA studies largely to locus-specific assays using sequencing or PCR based methods. Some studies have used methylation of long interspersed nucleotide element-1 (LINE-1) as a proxy measure of global DNA methylation level and observed hypomethylation in both tissue and plasma of CRC patients^15^. However, genome-wide studies including those with methylation array datasets for cfDNA remain scarce, limited to small sample numbers^16^ or pooled samples^17,18^. As a result, no *global* hypomethylation assay has translated into cfDNA diagnostics despite extensive tissue evidence. Yet global hypomethylation remains an appealing biomarker if it can be measured rapidly and inexpensively, as it offers a robust, heterogeneity-agnostic signal capable of segregating healthy and cancer samples at scale^19^.

Our group recently demonstrated a novel cfDNA assay called Asima Rev that analyzes the global structure of cfDNA - the “necklace” - rather than individual nucleotide-level events, by electrically measuring cfDNA aggregation as a surrogate biomarker for genome-wide methylation changes during cancer^20^. Global methylation changes are known to alter cfDNA’s physicochemical properties, including charge density, flexibility, conformation, and higher order structure, which together influence its electrical impedance signature. By measuring these properties in a rapid, label-less, and amplification-free manner on a chip, Asima Rev detects tumor-associated methylation shifts without the complexity of sequencing workflows and with potential pan-cancer applicability by capturing global methylation alterations conserved across malignancies.

In this study, we further evaluate the performance of Asima Rev for CRC detection and surveillance using archived plasma from treatment-naive CRC patients and age-matched controls. We assess performance across disease stages, explore the assay’s ability to detect recurrence, and investigate the biological basis of the assay signal. Specifically, we address: (i) whether global hypomethylation in cfDNA is a robust biomarker for CRC; (ii) whether Asima Rev truly measures global hypomethylation; (iii) how Asima Rev’s performance compares with existing CRC methylation assays; and (iv) the potential contribution of immune-derived cfDNA to the assay response. To contextualize Asima Rev’s biological signal, we analyzed 11 public methylation array datasets (**Table S1**), all generated using Illumina Infinium BeadChip arrays^21,22^, one of the most widely used high-throughput platforms for genome-scale methylation studies^8^. Methylation arrays have also been used in genome-scale studies in large projects such as The Cancer Genome Atlas (TCGA). Finally, we examined whether immune cells-derived cfDNA can confound Asima’s assay signals. cfDNA is derived from multiple tissues^23^, with blood cells being the dominant source in healthy individuals and cancer patients^23,24^, and tumor-derived cfDNA contributing additional fractions in cancer patients^25–27^. Given the limited availability of direct cfDNA methylome datasets, we evaluated immune-cell genomic DNA (gDNA) methylation patterns in CRC and in immune-related conditions. Our analyses suggest minimal influence of immune-derived DNA on the observed hypomethylation signals, though further validation is warranted.

## Results

### Diagnostic performance of Asima Rev

Diagnostic performance of Asima Rev assay was evaluated on 46 CRC patients and 33 healthy individuals (**Table S2**, **S3**). Impedance spectra of test samples were recorded across a wide frequency range (20 Hz to 30 MHz) to assess the aggregation differences between healthy and cancer-derived cfDNA (**Fig 1a**). cfDNA from healthy individuals consistently exhibited higher conductivity compared to cfDNA isolated from cancer patients (**Fig. 1b**), while the phase angle remained low and stable across all samples (**Fig. 1c**). Both Z and ΔZ values plateaued between 10 kHz and 1 MHz, with the phase angles in this region remaining < 2°, indicating dominantly resistive behavior. The maximum ΔZ was observed around 100 kHz (**Fig. 1d**), as previously reported by our group^20^. When ΔZ values were compared between healthy and CRC samples, a clear separation was observed (**Fig. 1e**), with stage I samples showing slightly higher mean values than more advanced stages (**Fig 1f**). Using ROC analysis, a binary classification threshold of 29 Ω was determined, yielding an AUC of 0.96 (**Fig. 1h**). At this threshold, the assay demonstrated an overall sensitivity of 95.65% (95% CI [85.47%, 99.23%]) and specificity of 93.94% (95% CI [80.39%, 98.92%]) for CRC detection. Sensitivity increased with disease stage as follows: 83.33% for stage I (95% CI [58.41%, 98.43%]), 100% for stage II (95% CI [78.47%, 100.0%]), 92.3% for stage III (95% CI [68.53%, 99.63%]) and 100% for stage IV (95% CI [78.47%, 100.0%]) (**Fig. 1g**). Among the CRC patients, six had history of cancer (see **Table S4**). Of these, five were correctly identified as positive by the assay, while one stage I CRC patient with prior leukemia was found to be false negative.

**Fig. 1.**
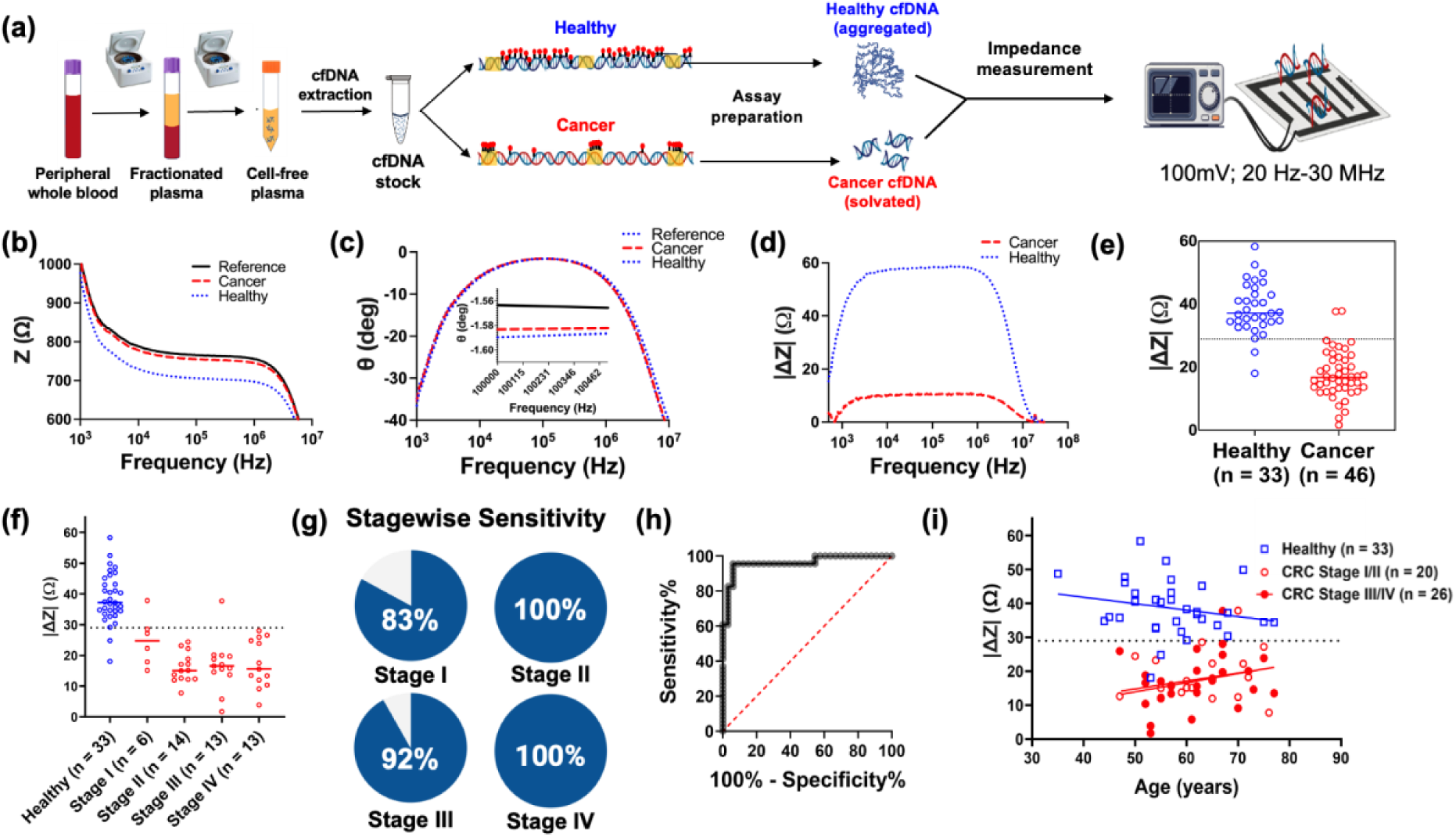
Performance of Asima Rev evaluated in CRC patients (n = 46) and healthy individuals (n = 33). **(a)** Schematic overview of the assay workflow; **(b-d)** Impedance spectra showing the magnitude (b) and phase angle of impedance (c), and the mod of differential impedance (|ΔZ|) (d) as a function of frequency for representative healthy (H19) and cancer (C21) samples. The inset in (c) shows the phase angle variation near 100 kHz; **(e, f)** Differential impedance values at 100 kHz across healthy (n = 33) and all CRC stages (n = 46) (e) and stagewise CRC patients (f). The dotted line indicates the diagnostic threshold at 29 Ω; **(g)** Pie chart showing stagewise sensitivity for CRC detection; **(h)** ROC analysis demonstrating high diagnostic performance, with an AUC of 0.96; and **(i)** Differential impedance as a function of age of healthy individuals, and early and late-stage CRC patients.

### CRC surveillance samples analyzed using Asima Rev

To explore the assay’s utility for CRC surveillance, we analyzed longitudinal plasma samples from six CRC patients (2 females, 4 males) collected over a 24-month period, pre- and post-treatment (**Fig. 2a**). All patients were treatment-naive, with no neoadjuvant systemic or radiation therapy, at baseline (t = 0) and had stage I-IV CRC (Table S2). Samples were included based on availability of serial collections with corresponding clinical data. Longitudinal |ΔZ| trajectories matched clinical status as documented in case report forms and corroborated by colonoscopy, imaging (CT, MRI), physical examinations, and pathology assessments. Where available, circulating carcinoembryonic antigen (CEA) levels were also included for comparison. In most cases, increasing |ΔZ| values corresponded to declining CEA levels and clinical remission, whereas decreasing |ΔZ| aligned with disease progression (**Fig. 2b-g**). For example, patient L4 showed rising |ΔZ| after a rectal carcinoid resection at visit 1, which later declined with the emergence of a new 4 mm lung nodule seen on follow-up CT highlighting the assay’s sensitivity to emerging metastatic lesions (Fig. 2e). Patient L1 showed increase in |ΔZ| after successful sigmoidectomy for removal of sigmoid colon adenocarcinoma at visit 1. Patient L1 was given adjuvant chemotherapy till visit 2 and had no clinically adverse reports at visit 2, followed by continued positive prognosis at visit 3, with continuously declining CEA levels (Fig. 2b). In patient L5, a slight decrease in |ΔZ| was observed (from 15 Ω to 10 Ω) during chemotherapy (5-Fluorouracil), despite tumor shrinkage (from 5.7 mm × 0.9 mm to 2.4 mm × 0.6 mm) as confirmed by MRI during visit 2. The |ΔZ| increased to 30 Ω, however, at visit 3 post treatment and with no radiologic evidence of disease.

**Fig. 2.**
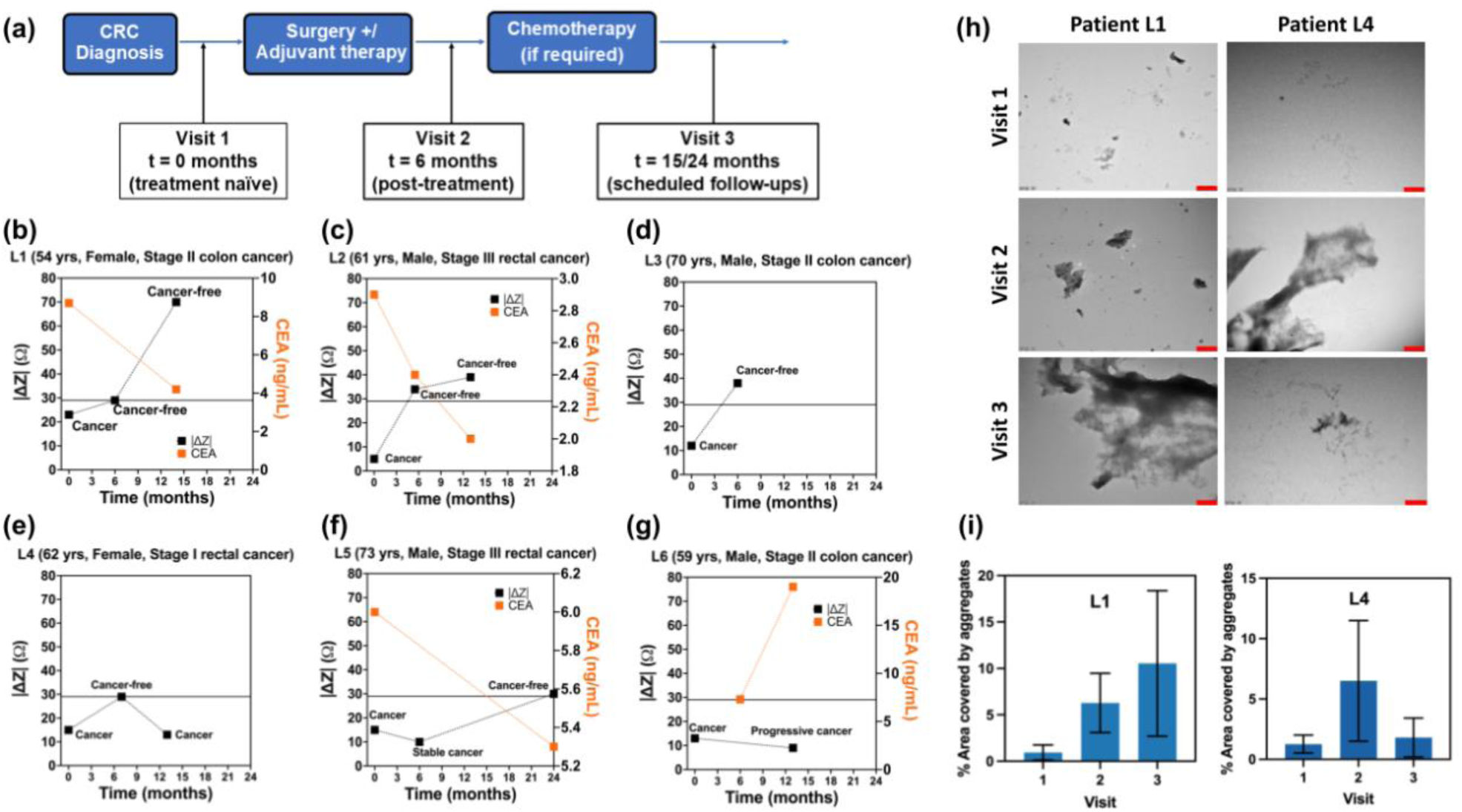
**(a)** Schematic showing timeline of longitudinal sample collection; **(b-g)** Longitudinal profiles of six CRC patients (L1-L6) over a 24-month period. Solid black data points represent |ΔZ| values at 100 kHz for each time point, with t = 0 denoting the treatment-naïve visit at confirmed diagnosis (clinical details for this visit are provided in the top legend on each graph). Cancer status at each time point is based on case report forms and clinical records. The solid black horizontal line indicates the diagnostic |ΔZ| threshold of 29 Ω. Solid orange data points indicate blood CEA levels, where available; **(h)** Representative TEM images of cfDNA samples from three different visits for patients L1 and L4. All the images were taken at 100 V and 15000x magnification. The scale bar is 500 nm; **(i)** TEM analysis across 90 images showing average area covered by the aggregates for visits 1, 2 and 3 for these two patients.

The discrepancy in |ΔZ| decrease in visit 2 for patient L5, despite tumor regression, may reflect lag in cfDNA clearance or other physiological influences such as biological aging, a known driver of global hypomethylation^28^. In general, our assay captured |ΔZ| variations despite normal age-related differences. **Fig. 1i** shows a negative association between ΔZ and age in healthy individuals (slope = −0.2495, p = 0.0013), contrasted by positive slopes in CRC patients (slope = 0.1357, p = 0.5170, and slope = 0.2690, p = 0.1739 for CRC stage I/II and CRC stage III/IV, respectively). Across the remaining longitudinal samples, the Asima Rev correctly reflected clinical remission or recurrence, though these patient-specific deviations highlight the need to investigate potential confounders affecting the |ΔZ| signal.

TEM analysis of samples from all three longitudinal visits of patients L1 and L4 confirmed cfDNA aggregation in samples, with the extent of aggregation correlating with the |ΔZ| values generated by Asima Rev. At visit 1, when both patients had confirmed diagnosis, TEM images showed minimal cfDNA aggregation (**Fig. 2h**), consistent with their low |ΔZ| values (23 Ω and 15 Ω) below the threshold. By visit 2, when both patients were clinically cancer-free, |ΔZ| values increased (Figs. 2b, 2e), and correspondingly, TEM revealed clear cfDNA aggregates. Patient L1 remained cancer-free at visit 3 and exhibited a high |ΔZ| of 70 Ω, much above the threshold, with TEM showing sustained high levels of aggregation. In contrast, patient L5 showed clinical recurrence at visit 3, accompanied by a decrease in |ΔZ| to 13 Ω (below the threshold) and a corresponding loss of visible cfDNA aggregation. While Fig. 2h provides representative images, all TEM fields captured in this study were quantitatively analyzed to determine average aggregate area, which aligned with previously reported methylation-dependent cfDNA structural changes (**Fig. 2i**). Together, these results support the mechanism that cfDNA morphology and charge distribution, modulated by methylation status, underpin the assay’s impedance-based detection mechanism.

### Estimating global DNA methylation levels from 450k array probes

Illumina 450k BeadChip array interrogates ∼450,000 CpG loci across the genome. After excluding missing, low-quality, cross-reactive, and gender-specific probes in the combined TCGA colon adenocarcinoma (COAD) and rectum adenocarcinoma (READ) datasets (**Fig. 3a**), average methylation across the remaining 398,783 probes (β*_av, all_*) was used to provide an initial estimate of global methylation differences between CRC tissue and adjacent normal tissue. Demographic and clinical characteristics of CRC patients from TCGA are provided in **Table S5**. The rationale for probe filtering is provided in the Supplementary Section (**Figs. S1, S2 and Note S1**). This “global” measure indicated modest hypomethylation in CRC tissue (β*_av, all_* = 0.4780) compared to adjacent normal tissue (β*_av, all_* = 0.4893) (**Fig. 3b**). The effect size was medium (Hedge’s g = 0.440) but with limited discriminative utility (ROC AUC = 0.66, 95% CI [0.61, 0.71] (**Fig. 3c**). Hypomethylation was observed across all CRC stages, with lowest β-values in stage IV tumors (**Fig. 3d**). The relationship between β*_av, all_* and age using linear regression analysis yielded no correlation in healthy tissue, and a positive correlation was found with age in both early-stage CRC (slope = 0.00043, p = 0.0049 for stage I/II) and late-stage CRC (slope = 0.00028, p = 0.0566 for stage III/IV) (**Fig. 3e**).

**Fig. 3.**
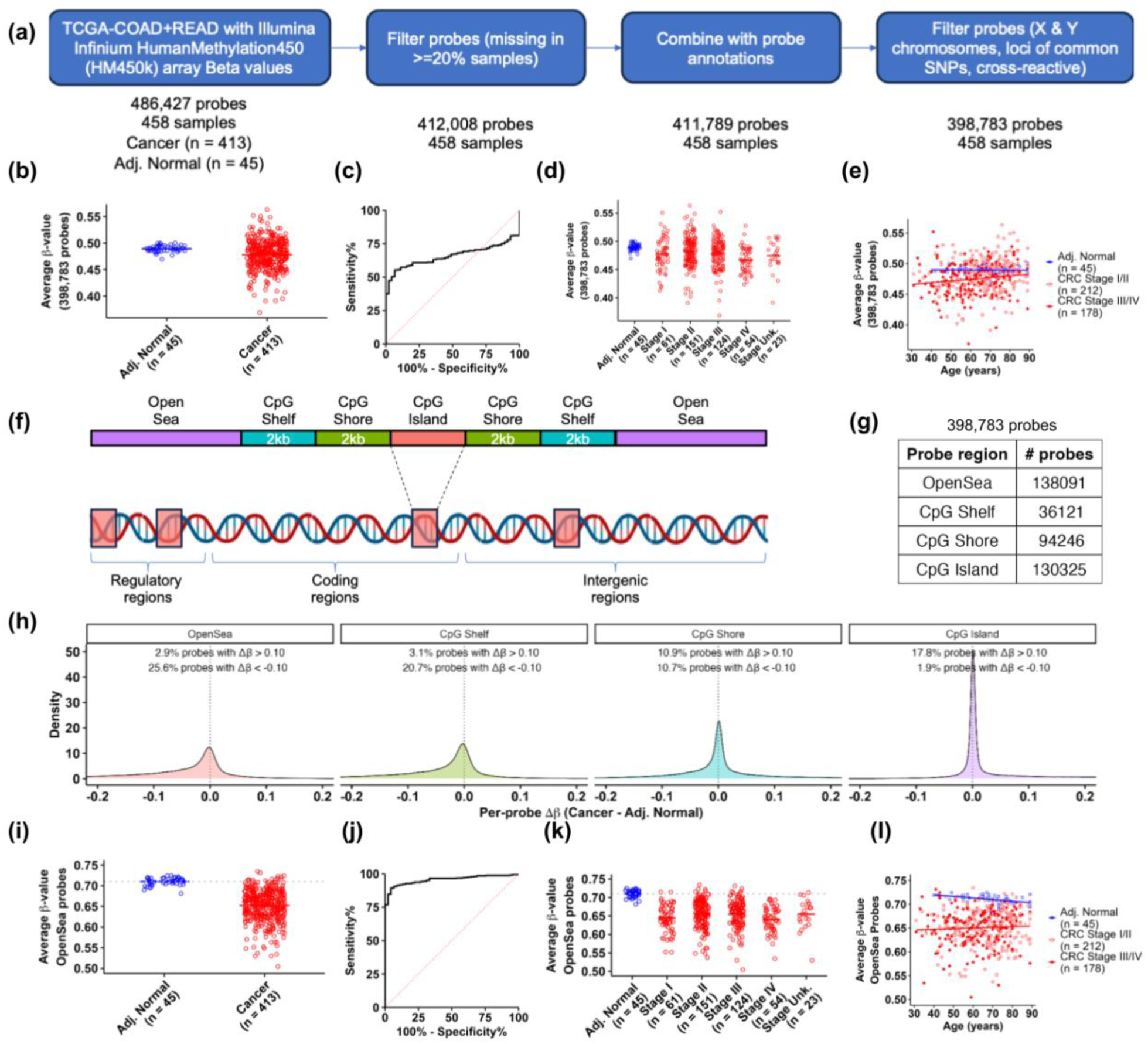
Average methylation of OpenSea probes captured global hypomethylation in CRC tissue. **(a)** An overview of methylation data preparation of TCGA CRC samples; **(b)** Comparison of average β-values across all 398,783 filtered probes (β*_av, all_*) in CRC and adjacent normal tissue samples; **(c)** ROC curve indicating low discriminatory ability (AUC = 0.66) for CRC detection; **(d)** β*_av, all_* for CRC patients separated by pathologic stage. Stage unk. refers to samples whose pathologic stage is not annotated in TCGA. **(e)** Linear correlation of β*_av, all_* with age in CRC samples for stage I/II, stage III/IV and adjacent normal tissue. **(f)** Schematic showing higher density of CpG islands in gene regulatory regions and their surrounding genomic contexts: CpG Shores (0–2 kb), CpG Shelves (2–4 kb), and OpenSea regions (> 4 kb); **(g)** Table showing number of probes belonging to each genomic region for the TCGA data. **(h)** Density of Δβ values (cancer - adjacent normal) for all probes belonging to each of the four genomic regions in (f); **(i)** Comparison of average β-value across all 138,091 OpenSea probes (β*_av, OS_*) in adjacent normal tissue and CRC. The dotted gray line shows β*_av, OS_* for adjacent normal tissue; **(j)** ROC curve indicating high discriminatory utility (AUC = 0.96) for CRC detection; **(k)** β*_av, OS_* for CRC patients separated by pathologic stage; **(l)** Linear correlation of β*_av, OS_* with age in CRC samples for stage I/II, stage III/IV and adjacent normal tissue.

The above estimates of genome-wide methylation were constrained by the intrinsic design bias of the Illumina 450k array, which includes probes targeting CpG islands, CpG Shores, CpG Shelves, and OpenSea regions of the human genome^21^ (**Fig. 3f**). By design, the array disproportionately targets CpG islands (30.9% of all probes^29^), which constitute only ∼0.7% of the genome, and under-represent OpenSea regions (36.3% of all probes^29^), which make up ∼93.3% of the genome^30^ (**Fig. 3g**). The percentage of hypomethylated (Δβ < −0.1) and hypermethylated (Δβ > 0.1) probes across different genomic regions revealed that only 1.9% of CpG island probes were hypomethylated in CRC, compared to 25.6% of all OpenSea probes (**Fig. 3h**). This imbalance led β*_av, all_* to substantially underestimate the extent of genome-wide hypomethylation due to over-representation of CpG island probes. Moreover, prior studies have indicated enrichment of OpenSea fragments in cfDNA^31^, further supporting their relevance for liquid biopsy assays. Accordingly, average β-value across OpenSea probes (β*_av, OS_*) provided a much more accurate and clinically meaningful measure of global hypomethylation in CRC. Consistent with this, β*_av, OS_* showed a strong concordance (R² = 0.977) with average methylation of LINE-1 probes (β*_av, LINE-1_*) (**Fig. S3**), which are established markers of global hypomethylation, while β*_av, all_* showed a substantially weaker correlation (R² = 0.686) (**Fig. S4**).

Using this measure, CRC tissue samples were found to be strongly and significantly hypomethylated (β*_av, OS_* = 0.652) compared to adjacent normal tissue (β*_av, OS_* = 0.710, Wilcoxon rank-sum test, p < 2.2e-16) (**Fig. 3i**), yielding a clinically meaningful ROC AUC of 0.96, 95% CI [0.94, 0.98] (**Fig. 3j**). Although β*_av, OS_* showed greater variance in CRC tissue than adjacent normal tissue samples, the hypomethylation pattern was consistent across all cancer stages (**Fig. 3k**). Linear regression analysis of β*_av, OS_* with age showed significant age-related hypomethylation in adjacent normal tissue (slope = −0.00034, p = 0.0025), but no significant change with age in CRC tissue (slope = 0.00030, p = 0.15, and slope = 0.00013, p = 0.53 for CRC stage I/II and CRC stage III/IV, respectively) (**Fig. 3l)**.

### Estimating DNA methylation across known molecular subtypes of CRC

Next, we compared β*_av, OS_* across four independent molecular subtypes, namely CRC with MSI and MSS; CRC with CIMP high, CIMP low and CIMP negative status; CRC with subtypes defined by Guinney *et al.*^11^ based on gene expression clustering, CMS1, CMS2, CMS3, and CMS4; and CRC with and without BRAF mutation. Compared to adjacent normal tissue (β*_av, OS_* = 0.710), the average methylation level in the MSS group was 0.649 (Wilcoxon rank-sum test, W = 347, p < 2.2e-16) and that in the MSI group was 0.669 (Wilcoxon rank-sum test, W = 325, p = 7.22e-11) (**Fig. 4a**). For CIMP, where differences exist in hypermethylation on CpG islands, the average global methylation level of CIMP high (β*_av, OS_* = 0.658) was similar to that of CIMP low (β*_av, OS_* = 0.650) and CIMP negative (β*_av, OS_* = 0.655), and all were significantly lower than the adjacent normal tissue (Wilcoxon rank-sum test, W = 176, p = 1.51e-12 for CIMP high, W = 67, p = 3.60e-16 for CIMP low, and W = 299, p < 2.2e-16 for CIMP negative) (**Fig. 4b**). Across CMS subtypes, we observed that the average methylation across CMS1 (β*_av, OS_* = 0.670) and CMS4 (β*_av, OS_* = 0.665) subtypes was slightly higher compared to CMS2 (β*_av, OS_* = 0.644) and CMS3 (β*_av, OS_* = 0.638). Each of these subtypes were, however, significantly hypomethylated compared to healthy adjacent tissue (Wilcoxon rank-sum test, W = 232, p = 1.13e-10 for CMS1, W = 114, p < 2.2e-16 for CMS2, W = 46, p = 1.88e-15 for CMS3, and W = 340, p < 2.2e-16 for CMS4) (**Fig. 4c**). In CRC patients with BRAF mutation, the average methylation level was 0.674 compared to BRAF wild type patients at 0.651, and both were significantly hypomethylated compared to adjacent normal tissue (Wilcoxon rank-sum test, W = 192, p = 4.26e-9 for BRAF Mut, and W = 458, p < 2.2e-16 for BRAF WT) (**Fig. 4d**). Overall, we found that despite minor differences in β*_av, OS_* between molecular subtypes in CRC patients, all subtypes were significantly hypomethylated compared to normal controls. These results suggested that in the absence of additional factors, we would expect assays based on global hypomethylation to work well across all CRC molecular subtypes. In comparison, β*_av, all_* values were sensitive to hypermethylation in CpG islands and underestimated global hypomethylation across CRC molecular subtypes (**Fig. S5**).

**Fig. 4.**
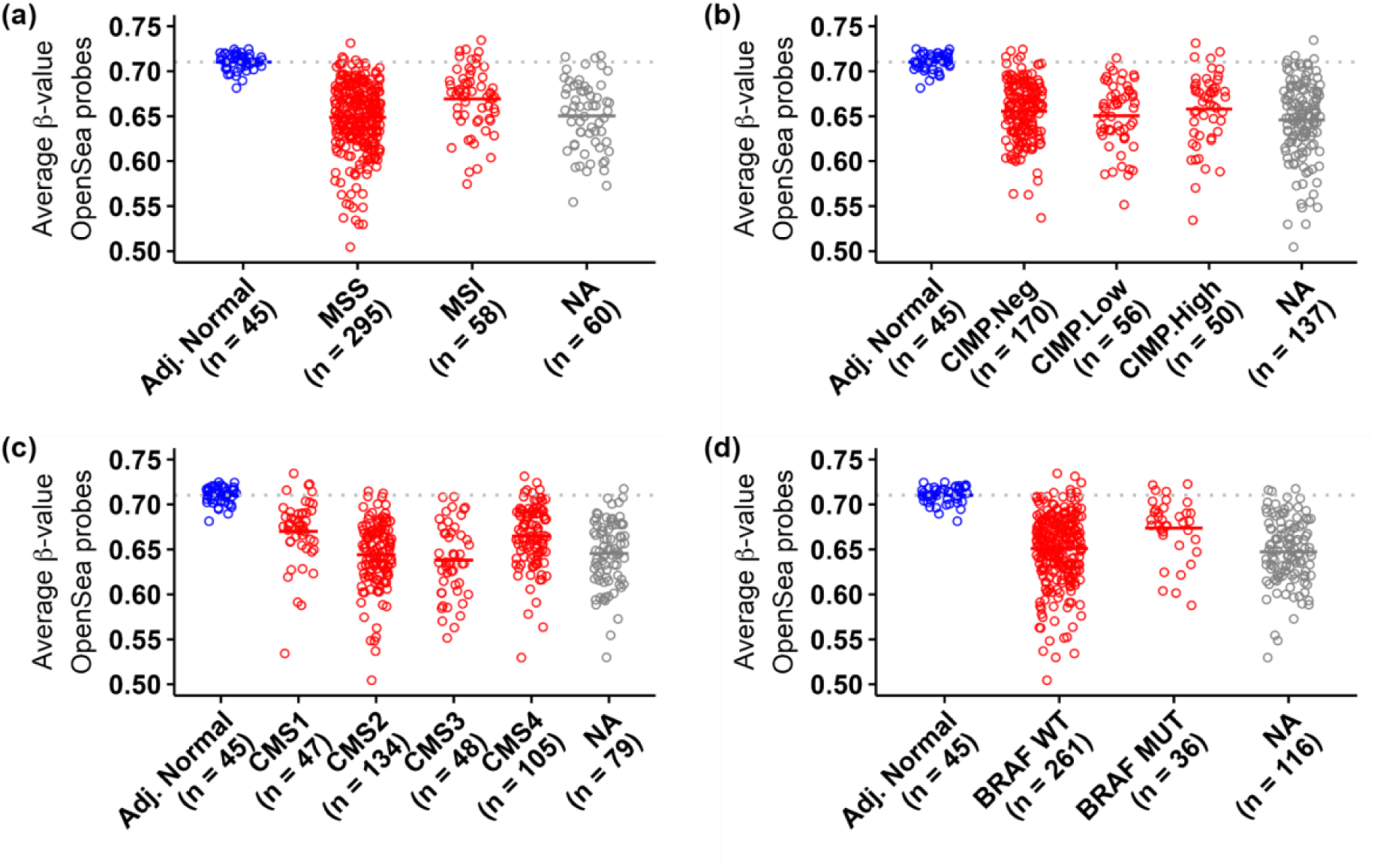
All CRC molecular subtypes showed hypomethylation compared to healthy controls in TCGA. Average β-value across OpenSea probes (β*_av, OS_*) in CRC patients with **(a)** microsatellite stability (MSS) and microsatellite instability (MSI); **(b)** CpG Island Methylator Phenotypes - High (CIMP.High), Low (CIMP.Low) and Negative (CIMP.Neg); **(c)** Consensus Molecular Subtypes (CMS) 1-4, and; **(d)** wild type (BRAF WT) and mutated (BRAF MUT) BRAF gene compared to adjacent normal tissue (Adj. Normal). NA refers to samples for which molecular subtype annotation was unavailable.

### Comparing DNA methylation in healthy individuals and adjacent normal tissue from CRC patients

To compare DNA methylation profiles between healthy individuals and adjacent normal tissue from CRC patients, we analyzed four public datasets. Although absolute β*_av, OS_* values for healthy individuals varied across datasets (0.670 in GSE131013, 0.669 in GSE42752, 0.747 in GSE48684, 0.575 in GSE199057), the methylation levels of healthy tissue from cancer-free individuals closely matched those of adjacent normal tissue from CRC patients within each study and were consistently higher than those of CRC tissue in all except one study (**Figs. 5a** & **S6-S8**). For example, in GSE131013, β*_av, OS_* was 0.670 in healthy individuals, 0.674 in adjacent normal tissue and 0.632 in CRC tissue (**Fig. 5a**).

**Fig. 5.**
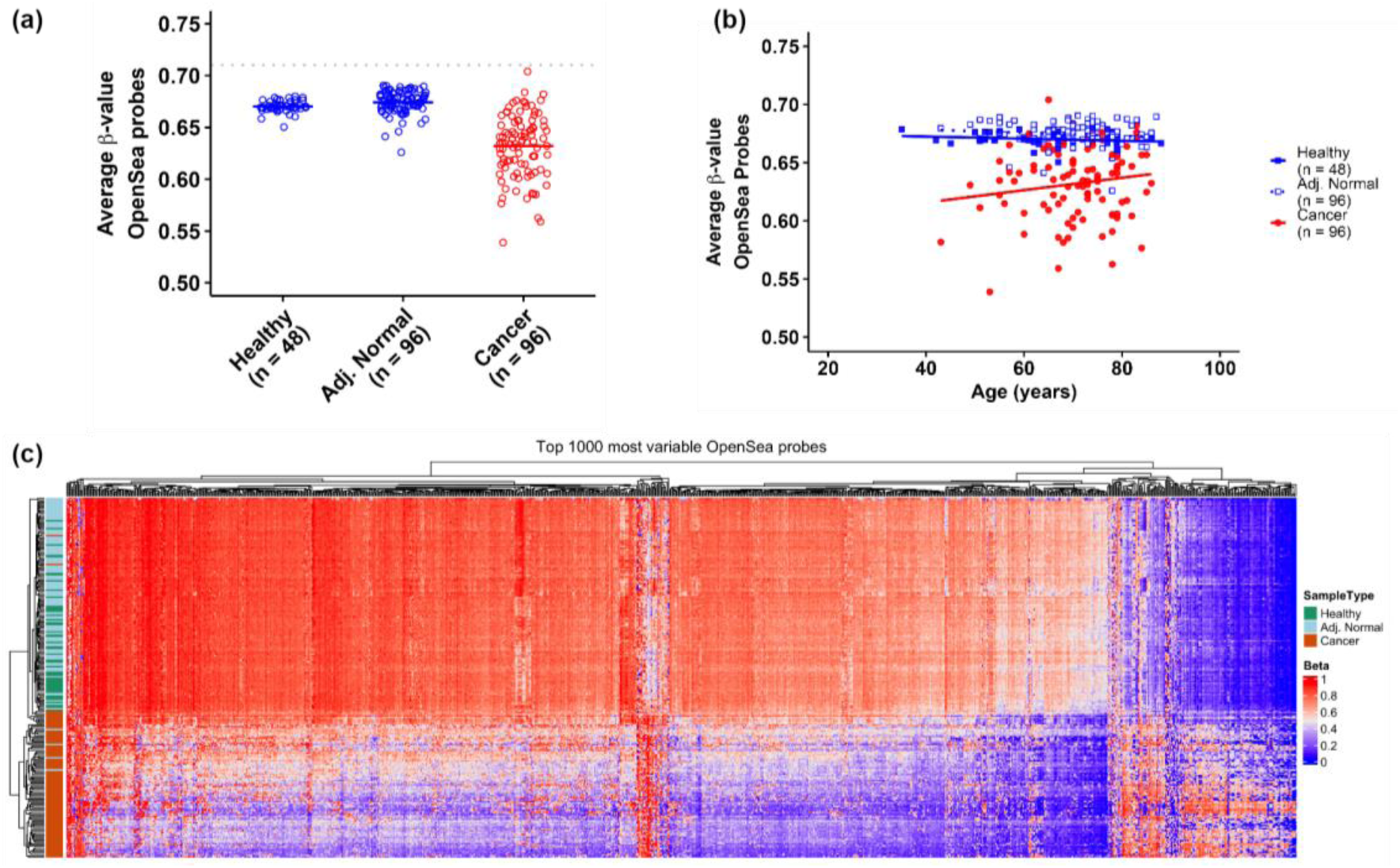
Average methylation in healthy individuals was similar to average methylation in adjacent normal tissue from CRC patients in GSE131013. **(a)** Comparison of average β-values across OpenSea probes (β*_av, OS_*) in healthy individuals (Healthy), adjacent normal tissue (Adj. Normal) from CRC patients, and CRC tissue (Cancer). The gray line represents β*_av, OS_* in adjacent normal tissue from the TCGA study. **(b)** Linear correlation of β*_av, OS_*with age in healthy individuals, adjacent normal tissue, and CRC samples; **(c)** Clustering analysis of the top 1000 most variable probes for healthy individuals, adjacent normal tissue from CRC patients, and CRC tissue.

Analysis of β*_av, OS_* with age indicated minimal but similar hypomethylation with age in healthy (slope = −0.00009, p = 0.196) and adjacent normal (slope = −0.00013, p = 0.261) samples, and an increase in methylation with age for CRC samples (slope = 0.0005, p = 0.13), although the trends were not statistically significant for any group (**Fig. 5b**). Clustering analysis of the top 1000 most variable OpenSea probes for all samples showed that the healthy and adjacent normal samples clustered together, indicating that these samples had similar methylation values locally at the level of individual probes, and not just at a global level (**Fig. 5c**). These findings support the use of adjacent normal tissue in TCGA (**Fig. 3**) as a suitable proxy for the methylation state of healthy individuals.

### Estimating global methylation in blood genomic DNA of CRC and autoimmune disease patients

We analyzed β*_av, OS_* in blood-derived gDNA from CRC patients and individuals with auto-immune diseases such as multiple sclerosis (MS) and rheumatoid arthritis (RA) to assess whether genomic hypomethylation in blood cells could interfere with Asima’s assay under different clinical conditions. In peripheral blood mononuclear cells (PBMCs) from CRC patients and healthy individuals, methylation levels were nearly identical (0.755 in healthy *vs.* 0.754 in CRC; Wilcoxon rank sum test: W = 1150, p = 0.491) (**Fig. 6a**). Similarly, in methylation array data from MS patients, average methylation in whole blood was comparable between healthy individuals (β*_av, OS_* = 0.737) and MS patients (β*_av, OS_* = 0.740; Wilcoxon rank-sum test, W = 106, p = 0.467) (**Fig. 6b**). Monocytes and WBCs from healthy individuals showed slightly lower methylation (0.724 and 0.722, respectively) than whole blood from either healthy or MS individuals. In peripheral blood leucocytes (PBLCs) from RA patients, the difference between healthy (β*_av, OS_* = 0.685) and RA samples (β*_av, OS_* = 0.682) was statistically significant (Wilcoxon rank-sum test, W = 50956, p =0.0014), but likely clinically indistinguishable (**Fig. 6c**). Collectively, these findings suggest that the hypomethylation-associated DNA aggregation captured by Asima’s cfDNA assay is more likely to be driven by tissue-specific hypomethylation in CRC rather than by methylation changes in blood cells. Whether cfDNA methylation differs between patients with autoimmune diseases and healthy individuals remains to be determined and is currently under investigation using Asima’s assay.

**Fig. 6.**
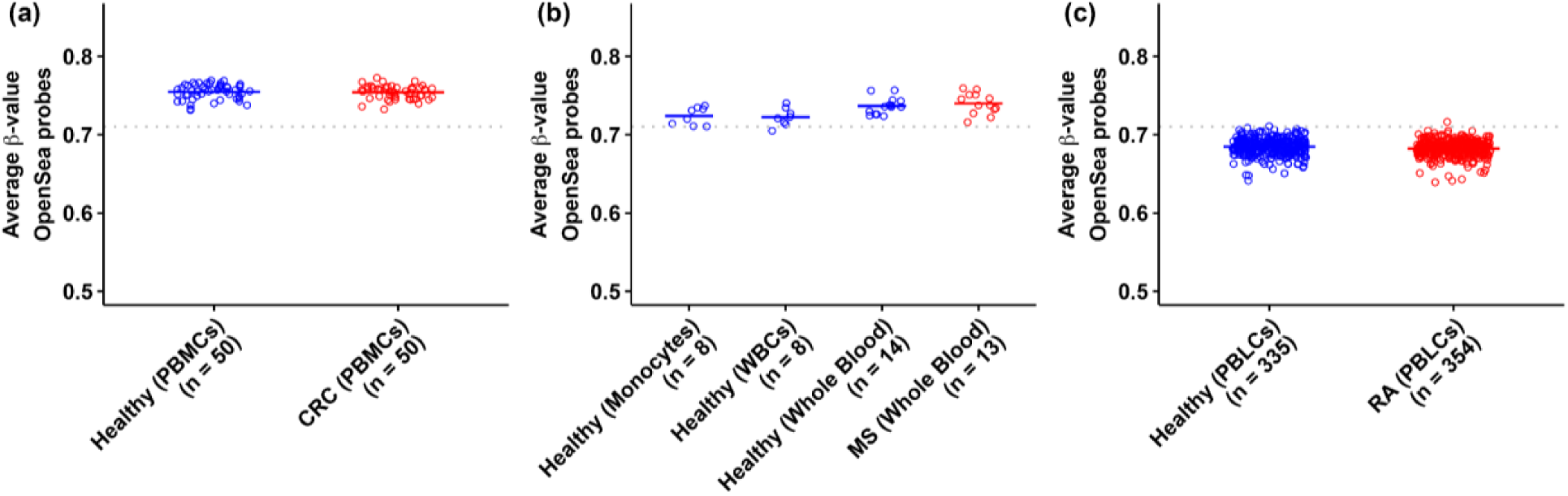
gDNA from blood samples of CRC and autoimmune disease patients did not show hypomethylation compared to healthy individuals. **(a)** Average β-values across OpenSea probes (β*_av, OS_*) in gDNA from peripheral blood mononuclear cells (PBMCs) in healthy individuals and CRC patients; **(b)** β*_av, OS_* in whole blood gDNA of MS patients and healthy individuals, and gDNA from monocytes and white blood cells (WBCs) from healthy individuals; **(c)** β*_av, OS_* in gDNA from peripheral blood leucocyte cells (PBLCs) in healthy individuals and rheumatoid arthritis (RA) patients. The gray line in each plot represents β*_av, OS_* in adjacent normal tissue from the TCGA study.

### Estimating global methylation in cfDNA samples from CRC patients

After establishing that OpenSea probes from Illumina 450k arrays capture global hypomethylation in gDNA from CRC tissues, we wanted to examine the same in cfDNA of CRC patients. In practice, Illumina 450k arrays require ∼250 ng DNA per test, which is difficult to obtain for cfDNA samples. Therefore, the number of methylation array studies on cfDNA is limited. We found only three array studies using cfDNA from CRC patients, including two studies, GSE110185 and GSE186381, from the same research group that used pooled samples. The third cfDNA study, GSE122126, did not use pooled samples but had small sample sizes with cfDNA from four CRC patients and four healthy individuals, and gDNA from colon epithelial cells of three healthy individuals. In this study, percentages of hypomethylated (Δβ < −0.1) and hypermethylated (Δβ > 0.1) probes in CRC cfDNA compared to healthy cfDNA across different genomic regions showed 3.6% of CpG island probes to be hypomethylated, compared to 24.8% of all OpenSea probes (**Fig. 7a**). These percentages were similar to those obtained in gDNA from TCGA samples, where 1.9% of all CpG island probes and 25.6% of all OpenSea probes were hypomethylated (**Fig. 3h**). The CRC cfDNA samples were also globally hypomethylated (β*_av, OS_* = 0.641) compared to healthy cfDNA (β*_av, OS_* = 0.676), and healthy gDNA from colon epithelial cells (β*_av, OS_* = 0.672) (**Fig. 7b**). We also examined the average methylation levels across LINE-1 probes and found CRC cfDNA samples to be hypomethylated (β*_av, LINE-1_* = 0.662) compared to healthy cfDNA (β*_av, LINE-1_* = 0.702) and gDNA from colon epithelial cells (β*_av, LINE-1_* = 0.715) (**Fig. 7c**). The overall trend of global hypomethylation in this study was similar to that of gDNA from CRC tissue samples, but the number of samples was too small to draw statistical conclusions. The two studies with pooled cfDNA samples, did not show average methylation difference between pooled samples from healthy individuals and CRC patients (**Figs. S9, S10**).

**Fig. 7.**
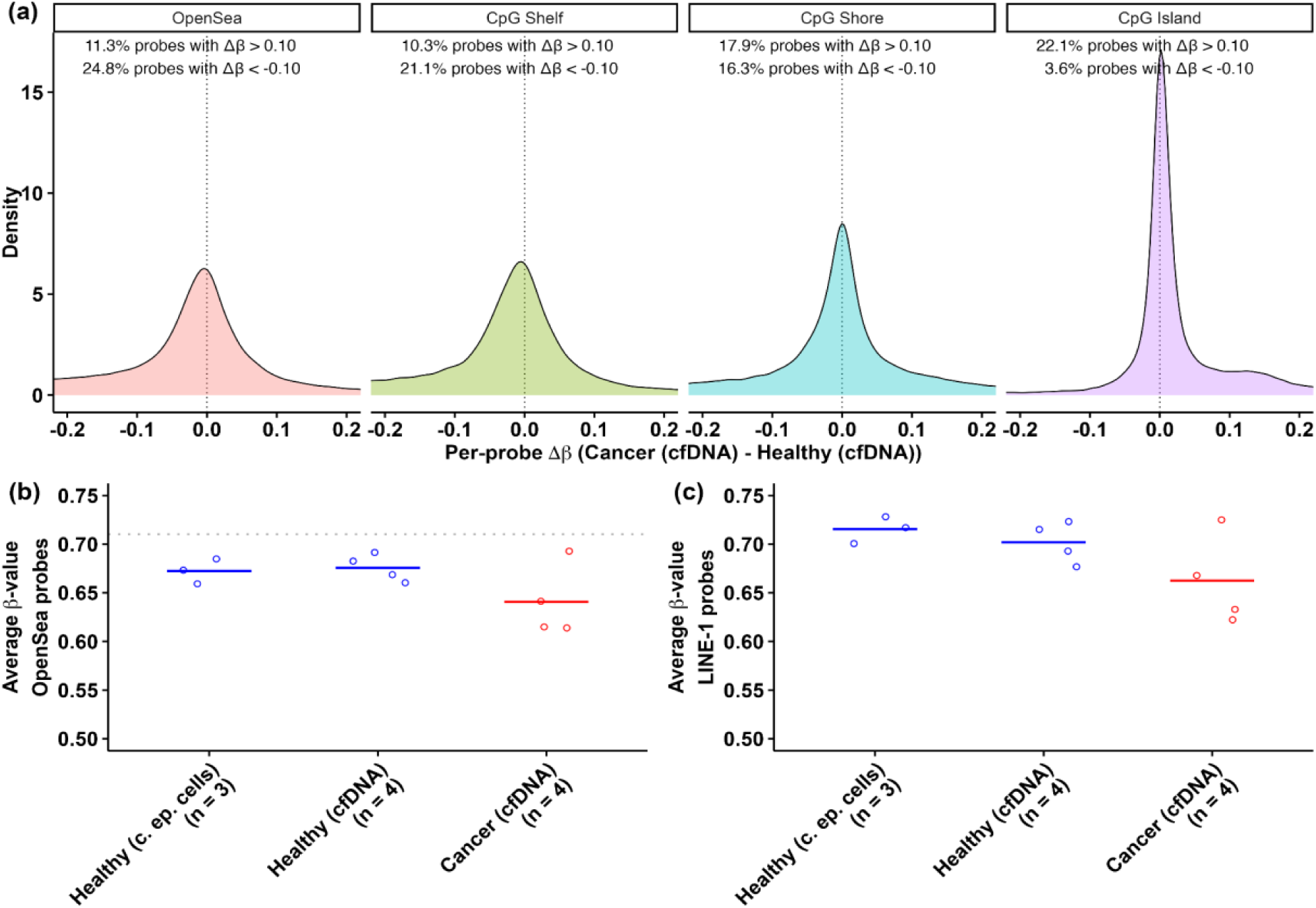
Average methylation of OpenSea probes in cfDNA showed global hypomethylation in CRC patients in GSE122126. **(a)** Density of Δβ values for all probes belonging to each of the four genomic regions on 450k array: OpenSea, CpG Shelf, CpG Shore, and CpG island; **(b)** Comparison of average β-value across OpenSea probes (β*_av, OS_*) in gDNA from colon epithelial cells (c. ep. cells), cfDNA from healthy individuals and cfDNA from CRC patients; **(c)** Comparison of average β-value across LINE-1 probes (β*_av, LINE-1_*) in gDNA from colon epithelial cells, cfDNA from healthy individuals and cfDNA from CRC patients

## Discussion

In this study, we demonstrated that measuring cfDNA aggregation in plasma can reliably discriminate between healthy individuals and patients with CRC, and that the same signal tracks with cancer recurrence in patients following curative treatment. Our underlying hypothesis is that the electrical signal differences measured by Asima Rev are driven by methylation-dependent aggregation in cfDNA. cfDNA from healthy individuals, which contains a higher number of methylated cytosines, tends to self-assemble into microaggregates in aqueous environments due to the hydrophobic nature of the methyl group. As methyl cytosines are progressively lost during cancer development, cfDNA transitions from microaggregated structures to a nanoaggregated state. The observed reduction in electrical signal in the Asima Rev assay was corroborated with TEM results showing a reduction in aggregation. Building on this observation, we analyzed available public methylation data sets to further characterize global methylation status of gDNA from cancer and adjacent tissues, and from plasma cfDNA to corroborate the potential DNA hypomethylation as a key biomarker for cancer assessment. Our analysis confirmed that global hypomethylation is consistently detectable in CRC tissue, particularly when examining OpenSea regions.

Methylation array datasets are predominantly used to study gDNA. Prior studies indicate that cfDNA is enriched for OpenSea fragments^31^ - regions underrepresented on methylation arrays, which are strongly biased toward CpG islands (∼0.7% of the genome) and gene promoters^21,22^. To address this mismatch, we restricted array analyses to OpenSea probes, aligning coverage more closely with cfDNA composition. This approach is consistent with studies using repetitive elements such as LINE-1 as global methylation proxies^32^. After truncation of the probe dataset, we observed strong concordance between Asima rev results in cfDNA and global hypomethylation in tissue samples (specificity 95.56% (95% CI [88.89, 100]) and sensitivity 89.35% (95% CI [86.2, 92.3]) for TCGA cohort), supporting the conclusion that Asima Rev reflects true genome-wide methylation loss. Notably, Asima Rev’s performance exceeded that of conventional locus-specific assays, presumably because it captures structural consequences of truly global, rather than site-restricted, hypomethylation.

A central question raised by our work is why the robust hypomethylation, so consistently observed in CRC tissue, yields inconsistent findings across the limited cfDNA methylation literature. Our analysis identified significant global hypomethylation in only one published cfDNA methylation array study but not in others. Several technical and biological factors likely contribute to this heterogeneity. Methylation arrays require relatively large DNA input (∼250 ng), often necessitating investigators to pool cfDNA samples, potentially introducing bias or diluting tumor-derived signals. The cfDNA studies we analyzed varied in sample preparation protocols, pooling strategies, and cohort characteristics, all of which may influence detection sensitivity. Moss *et al.*^23^ reported observable hypomethylation in individual cfDNA samples from CRC patients, whereas Gallardo-Gomez *et al.*^17,18^ used pooled samples where global hypomethylation signal may have been diluted or lost. Tumor fraction in cfDNA is known to be highly variable, with circulating tumor DNA (ctDNA) estimated in the range of 0.1 - >10% of the total cfDNA^25^. Array-based methods, which measure average methylation across bulk cfDNA, may lack the sensitivity to detect subtle global shifts when tumor-derived DNA is sparse. Moreover, biological heterogeneity in cfDNA composition, reflecting contributions from tumor, stroma, immune cells, and normal tissue turnover^26,27^, adds complexity. Tumor-derived cfDNA in CRC patients exhibits characteristic fragmentation patterns^33^, chromatin accessibility signatures^34^, and nucleosome positioning^35^, all of which may modulate aggregation behavior independently of or synergistically with methylation status. Our observation that immune cell gDNA does not exhibit hypomethylation in CRC patients or autoimmune conditions suggests that changes in immune-derived cfDNA are unlikely to confound the Asima Rev signal, though sequencing-based analysis of immune cells and cfDNA will be needed for definitive answers (**Fig. 8**).

**Fig. 8.**
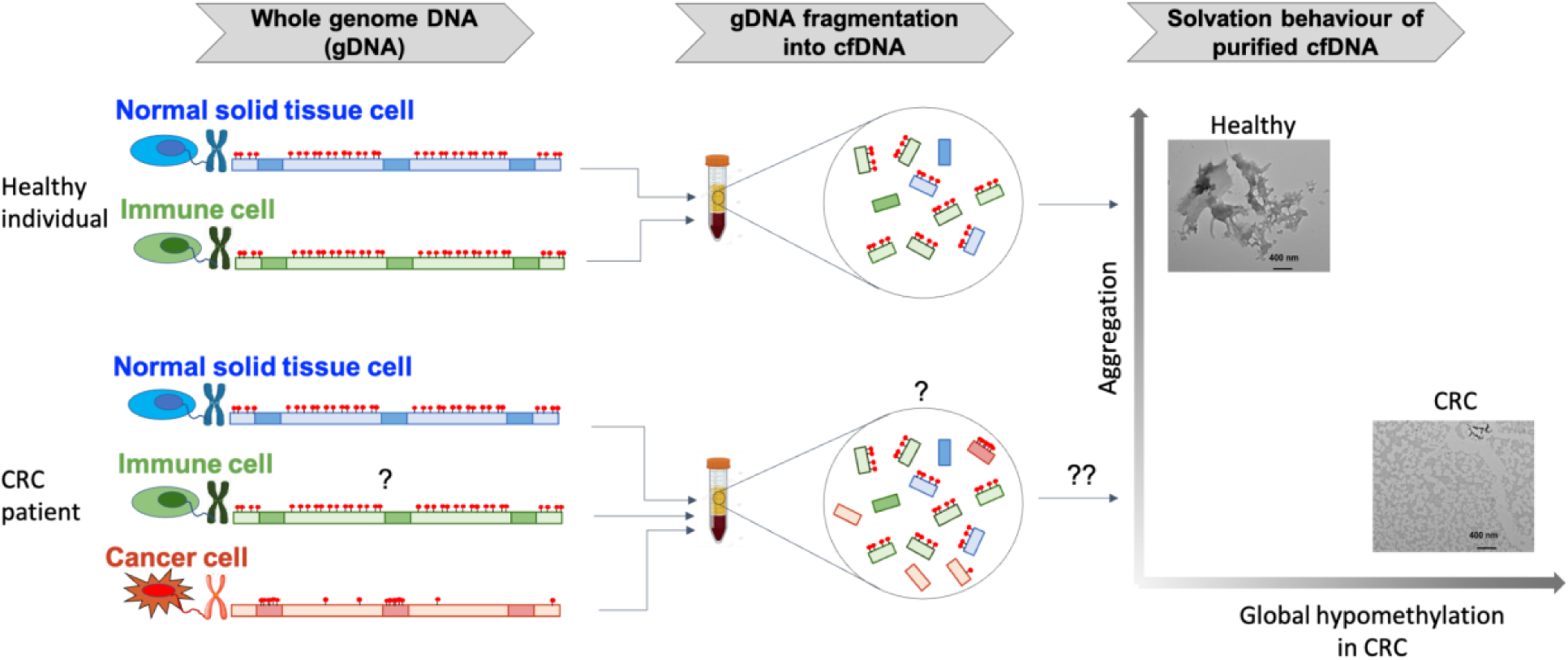
Schematic overview of the origin, proposed fragmentation and observed solvation behaviour of cfDNA in healthy individuals and CRC patients.

While we have not definitively established global hypomethylation in cfDNA of CRC patients in this study, we have established different aggregation behaviour in healthy and cancer cfDNA. Since Asima Rev measures aggregation behavior, which is a collective biophysical property, the assay may amplify even small shifts in methylation status through cooperative interactions among cfDNA molecules. This phenomenon parallels observations in DNA condensation and chromatin structure, where small changes in charge density can produce disproportionately large effects on higher-order organization. It is also possible that Asima Rev integrates multiple features beyond methylation alone. cfDNA fragment length, end motifs, and nucleosome occupancy are all altered in cancer^19^ and may contribute to aggregation dynamics and provide robustness against tumor heterogeneity.

Several limitations merit consideration. First, while we demonstrate strong correlation between Asima Rev signals and tissue-based global hypomethylation, direct bisulfite sequencing of cfDNA from assayed samples was not performed in this study. Second, we focused only on global hypomethylation; integration of fragmentomics, chromatin accessibility, histone post-translational modifications on nucleosomes^36^ or nucleosome positioning data could further elucidate the basis of aggregation behaviour. These phenomena accompany methylation changes and likely have significant impact on charge, hydrophobic/philic and conformational properties of the DNA. Third, confounding contributions from non-tumor cfDNA sources require further evaluation. While our analysis of immune cell gDNA suggests minimal methylation changes in CRC or autoimmune contexts, other inflammatory conditions, infections, and tissue injury were not considered. Autoimmune conditions such as MS and RA may also affect downstream cfDNA methylation, which was not evaluated in this study. Fourth, while we analyzed tissue gDNA, finding hypomethylation across all CRC molecular subtypes, we did not have access to cfDNA samples with different sub-types, and these will be evaluated in future work.

Finally, in our prior study, we showed that the Asima Rev assay could distinguish cancer from normal across more than 15 cancer types; however, only a small number of CRC samples were included (n = 4). The present study builds on those findings by demonstrating that the assay performs robustly for CRC detection as well, achieving high sensitivity and specificity. For context, the best-in-class FDA-approved blood-based cfDNA assay, Guardant Shield, reports 83% sensitivity and 90% specificity. Unlike tissue-specific assays, however, our test is pan-cancer in nature, and in a screening setting, would require a follow-up diagnostic test to establish tissue of origin.

### Conclusions

In this study, we demonstrated that Asima Rev, a rapid, label-free electrical impedance assay that measures cfDNA aggregation, achieves clinically meaningful performance for CRC detection. The assay distinguished CRC patients from healthy individuals with 95.65% sensitivity and 93.94% specificity across all cancer stages. In preliminary analyses, the assay also showed promise for longitudinal monitoring, however this requires validation in larger sample sizes and ideally in a prospective study. These findings establish Asima Rev as a clinically viable approach for CRC detection by leveraging global hypomethylation–driven changes in cfDNA aggregation.

While the precise molecular composition and methylation landscape of cfDNA remain active areas of investigation, our results provide compelling evidence that global hypomethylation is not only detectable in cfDNA but diagnostically actionable. Our assay supports a new paradigm in cfDNA-based cancer diagnostics, one that emphasizes structural profiling over base-by-base sequencing. Our findings from public methylation data bridge a gap between tissue-based observations of global hypomethylation and their translation to liquid biopsy applications for CRC, offering biological insight and generating new hypotheses to explore cancer epigenetics in cfDNA. As healthcare systems face mounting demands for accessible, cost-effective cancer screening, technologies that are simple, rapid, and accurate will be essential. The performance and operational simplicity of Asima Rev position it as a promising tool for early CRC detection and surveillance, with strong potential to increase screening uptake and expand access in underserved and resource-limited settings. Ongoing and future work will focus on mechanistic dissection of the relationship between methylation-dependent DNA structural changes and impedance behavior, including controlled perturbation experiments and orthogonal biophysical measurements.

## Materials and Methods

### Materials

Platinum-on-glass planar interdigitated electrodes (IDEs) with 40 symmetrical fingers (20 pairs) of 3.5 mm length and 100 μm width each, and 100 μm edge-to-edge spacing (Micrux Technologies, Spain); Sodium hydroxide (NaOH) (Sigma Adrich, Canada); MagMax cell-free DNA isolation kit (A29319, Thermofisher); 4-(2-hydroxyethyl)-1-piperazineethanesulfonic acid (HEPES) (Sigma-Aldrich, Canada); Polydimethylsiloxane (PDMS) sylgard 184 elastomer kit (Dow, Ellsworth Adhesives, Canada); K2/K3EDTA blood collection tubes (Greiner Bio-One, VWR Canada); 200 mesh square carbon TEM grids ( Electron Microscopy Sciences); UranyLess EM stain (Electron Microscopy sciences); Cryovials (Rose Scientific Ltd.). Buffer was prepared at pH 7.4 in ultrapure deionized Milli-Q water (∼18.2 MΩ.cm resistivity) following standard procedures and experiments were conducted at room temperature (RT ∼ 21 - 23 °C).

## Experimental Protocol

### Plasma collection

A total of 46 double-spun, cell-free frozen plasma vials were obtained from treatment-naïve patients with confirmed CRC, through Ontario Tumor Bank (Toronto, ON, Canada; n = 24), PrecisionMed BioIVT (Carlsbad, CA, USA; n = 18), and the Rajiv Gandhi Cancer Institute and Research Centre (Delhi, India; n = 4) (IRB-approved studies: 2023-3365-16254-6, 2023-3365-16271-4, and Res/BR/TRB-19/2020/56). The cohort included 24 males and 22 females, aged 45–75 years, encompassing all four CRC stages (stage I: 6; stage II: 14; stage III: 13; stage IV: 13) (see Table S2 for details). Additionally, longitudinal plasma samples collected over a 24-month period were available for six PrecisionMed patients. For controls, 33 blood samples were collected from healthy individuals (15 males, 18 females; aged 35–75 yrs) using the same protocol as the cancer samples and after obtaining informed consent (see Table S3 for details). Peripheral blood (6 mL) was drawn via venous phlebotomy into K2EDTA tubes. Plasma was processed within 4 h of collection (with blood exposure to RT limited to < 1 h). Initial centrifugation was performed at 2000g for 10 min at 4°C, followed by transfer to 15 mL Falcon tubes and a second centrifugation at 3000g for 15 min at 4°C. The resulting double-spun plasma was aliquoted into 1 mL cryovials and stored at –80°C until further processing.

### cfDNA extraction and quantification

Plasma samples were thawed by transferring them to −20°C overnight, followed by incubation at RT for 30 min. cfDNA was then extracted using the MagMax™ Cell-Free DNA Extraction Kit (A29319, Thermo Fisher Scientific), following a slightly modified protocol. Briefly, the plasma was mixed with the recommended volumes of lysis/binding solution and magnetic beads, then vortexed for 20 min. The mixture was placed on a magnetic stand for 3-5 min to separate the cfDNA-bound magnetic beads. Beads were washed twice with the wash solution and twice with 80% ethanol. After air drying, cfDNA was eluted in 20 μL of elution buffer, referred to as the “cfDNA stock.” cfDNA concentration in the stock (C_stock_) was quantified using both a NanoDrop One™ spectrophotometer and a Qubit™ 4 fluorometer (Thermo Fisher Scientific), following the manufacturers’ protocols.

### cfDNA sample preparation

One μL of cfDNA stock was added to 49 μL of 15 mM HEPES buffer (pH 7.4) and mixed vigorously. The sample was then stored at 4 °C for 4-5 h followed by equilibration at RT prior to impedance measurements. The cfDNA stock volume in HEPES was chosen such that the cfDNA concentration in the sample was identical to that in the plasma from which it was extracted. The plasma (or sample) cfDNA concentration was estimated using the following equation: C_plasma_ = (C_stock_ x V_stock_)/V_plasma,_ where C is the concentration and V is volume. A reference solution was also freshly prepared by adding 1 μL of elution buffer to 49 μL of HEPES buffer and used for baselining the impedance measurements.

### Impedance measurement

A PDMS microchamber (4 mm diameter) was placed over the IDEs. Twenty microliters of the reference solution were added to the chamber, which was then sealed with a coverslip to minimize evaporation. Impedance measurements were performed using an ISX-3 impedance analyzer (Sciospec Scientific Instruments GmbH, Germany) over a frequency range of 20 Hz to 30 MHz with an excitation voltage of 200 mVpp, following load compensation with a 1 kΩ resistor. The test sample was measured under identical conditions immediately after the reference. Total impedance (Z) and phase angle (θ) were calculated from the real (Z′) and imaginary (Z″) components. Final results were expressed as the absolute differential impedance values between the test and reference samples, |ΔZ| = |Z_test_ – Z_ref_|, and |Δθ| = |θ_test_ – θ_ref_| to reduce inter-sample variability. All downstream analyses were conducted at 100 kHz, based on our previous findings^20^. Wherever possible, duplicate samples were prepared independently from the same stock to check the reproducibility of the data points (see Tables S2, S3 for standard deviations). Receiver operating characteristic (ROC) curve analysis for sensitivity and specificity was performed using GraphPad Prism (version 10.3.1).

### TEM characterization

Prepared cfDNA samples were analyzed using transmission electron microscopy (TEM). Briefly, 10 *μ*l of the sample was dropped on a copper mesh coated TEM grid and blotted from the other side of the grid. The TEM grids are then stained with Uranyless EM stain for 1 min, followed by washing with distilled water. The grids were air-dried overnight and then imaged at different magnifications using TEM (Zeiss TEM Libra 120) at 100 kV. These images were then analyzed using ImageJ software (Fiji, version 2.16) to estimate the average of the percentage area covered by the aggregates, which was then plotted as an average ± 1SD (see **Table S6** for analysis details).

## Methylation analysis of Illumina 450k and EPIC arrays from public studies

### Data preparation from TCGA

Illumina 450K methylation array data for colon adenocarcinoma (COAD) and rectum adenocarcinoma (READ) were downloaded from TCGA via the GDC portal (https://portal.gdc.cancer.gov/). Metadata, including sample processing and clinical information, and pre-processed β-values, were obtained. The β-values, ranging from 0 to 1, represent the average methylation level for each of the 486,427 probes in every sample. The dataset contained 458 samples, including 413 tumor samples and 45 adjacent normal tissue controls. Each sample was annotated with age, sex, primary diagnosis and pathologic stage using associated metadata. Probes whose values were missing in more than 20% of the samples were removed prior to downstream analysis. Principal components analysis (PCA) was performed on the top 1000 most variable probes across all samples, and it was observed that the adjacent normal samples clustered based on gender (**Fig. S1**). Therefore, probes on X and Y chromosomes were filtered out, along with the probes that are known to be cross-reactive^37^ and those that fall within commonly known Single nucleotide polymorphisms (SNPs). The remaining 398,783 probes were used for further analysis (**Fig. S2, Table S7**).

### Data preparation from Gene Expression Omnibus (GEO)

Pre-processed *β*-values were downloaded from GEO using the GEOquery package^38^ from Bioconductor. In some studies, *β*-values for each probe were available along with p-values. In these cases, only probes with a p-value of less than 0.05 in each sample were considered for analysis. Filtering was applied for probes missing in more than 20% of the samples in each study, probes on X and Y chromosomes, probes in commonly known SNPs and probes that are known to be cross-reactive, as described in the TCGA data section. In four studies that used 850k arrays, only probes that belong to 450k arrays were retained. The total number of probes available from source, number of probes with 450k array annotations, number of probes remaining after applying all filtering steps, and number of OpenSea probes in all studies are provided in **Table S7**.

### Annotation of genomic regions

Genomic regions for each probe, such as the chromosome location and classification of each probe as belonging to CpG island, CpG shore, CpG shelf or OpenSea region were annotated using the IlluminaHumanMethylation450kanno.ilmn12.hg19^39^ package from Bioconductor for all studies.

### Annotation of molecular subtypes in TCGA data

TCGA samples were annotated into four categories based on gene expression data - CMS1, CMS2, CMS3, and CMS4, two categories based on MSI and MSS status, three categories based on CIMP high, low, and negative subtypes, and BRAF mutation status based on a file released by Colorectal Cancer Subtyping Consortium^11^, available at https://doi.org/10.7303/syn2623706.

### Annotation of LINE-1 probes

Probes belonging to LINE-1 regions were annotated using the AnnotationHub package^40^ from Bioconductor. 450k probe coordinates (hg19) were obtained from the IlluminaHumanMethylation450kanno.ilmn12.hg19 package^39^ and converted to genomic ranges. RepeatMasker annotations were downloaded and filtered to LINE-1 elements (repClass = “LINE”, repFamily = “L1”). Probe overlaps with LINE-1 intervals were identified with *findOverlaps* function from GenomicRanges package^41^, and probes with at least one overlap were retained as LINE-1–associated CpGs.

### Data quality checks

Data quality checks were performed using the methylation status of probes associated with five genes described by Lu *et al.*^42^. The methylation level of probes associated with N4BP2 (cg13107169), EGFL8 (cg16282679), CTRB1 (cg16863382), TSPAN3 (cg21377793), and ZNF690 (cg12784172) was found consistently high across thirteen human tissue types and twenty-four cell lines. Therefore, these probes were considered similar to “housekeeping genes” in gene expression data analysis and used for quality checks on methylation array datasets.

### Comparison of methylation between samples

Three different proxies for the average methylation of a sample in a study were calculated - average *β*-value of all probes after filtration (β*_av, all_*), average *β*-value of all probes annotated as OpenSea (β*_av, OS_*), and average *β*-value of all probes annotated as LINE-1 (β*_av, LINE-1_*). The metric used was specified in each figure and the corresponding text. For quantification of hypermethylated and hypomethylated probes in each genomic region, Δβ was calculated by subtracting the average *β*-value of each probe in healthy or adjacent normal samples from the average *β*-value in cancer samples. Probes with Δβ < −0.1 were classified as hypomethylated and those with Δβ > 0.1 as hypermethylated. The percentage of probes meeting these criteria in each region was reported.

### Statistical analyses

All statistical analyses and visualizations were performed using R (v4.5.1) and Bioconductor packages unless otherwise noted.

## Acknowledgements

SG is grateful to Dr. Megan Hitchins, Dr. Theo deVos and Dr. John Abran for reviewing this manuscript. SK is grateful to Mikaeel Abbas for literature review.

