## Supplementary File for "Global Hypomethylation in Cell-free DNA Enables Non-invasive Colorectal Cancer Screening: Results from a Retrospective Validation Study"

### *Supplementary Information*

**Table S1.** The list of public methylation datasets analyzed in this study.

| No. | Data Source | Data Description | Sample Source | No. of Samples | Platform |
|---|---|---|---|---|---|
| 1 | TCGA-COAD and TCGA-READ | CRC patients | Tissue gDNA | 458 | 450k |
| 2 | GSE131013 | Healthy individuals and CRC patients | Tissue gDNA | 240 | 450k |
| 3 | GSE42752 | Healthy individuals and CRC patients | Tissue gDNA | 63 | 450k |
| 4 | GSE48684 | Healthy individuals and CRC patients | Tissue gDNA | 147 | 450k |
| 5 | GSE199057 | Healthy individuals and CRC patients | Tissue gDNA | 229 | 850k |
| 6 | GSE240324 | Healthy individuals and CRC patients | PBMC gDNA | 100 | 850k |
| 7 | GSE88824 | Healthy individuals and Multiple Sclerosis patients | Whole blood gDNA | 83 | 450k |
| 8 | GSE42861 | Healthy individuals and Rheumatoid Arthiritis patients | PBLC gDNA | 689 | 450k |
| 9 | GSE110185 | Healthy individuals and CRC patients | Whole blood pooled cfDNA | 6 | 850k |
| 10 | GSE186381 | Healthy individuals and CRC patients | Whole blood pooled cfDNA | 28 | 850k |
| 11 | GSE122126 | Healthy individuals and CRC patients | Whole Blood cfDNA | 7 | 450k |

**Table S2.** Clinical information of CRC patient samples used for cfDNA extraction. Information where not available is left blank.

| Sample ID | Cancer stage | Site of primary cancer | Age (yrs) | Sex | cfDNA stock conc. measured using Nanodrop (ng/µL) | cfDNA stock conc. measured using Qubit (ng/µL) | $\lvert\Delta Z\rvert$ at 100 kHz (ohms) | $\lvert\Delta\theta\rvert$ at 100 kHz (deg) |
|---|---|---|---|---|---|---|---|---|
| C1 | III | Colon | 65-69 | F | 2.2 | 0.354 | 19.71 ± 0.71 | 0.02 |
| C2 | IV | Colon | 45-49 | F | 2.5 | 0.434 | 25.95 ± 2.83 | 0.00 |
| C3 | IV | Colon | 65-69 | M | 4.2 | 0.583 | 24.84 ± 2.12 | 0.01 |
| C4 | III | Colon | 70-74 | M | 3.1 | 0.206 | 20.04 ± 7.07 | 0.04 |
| C5 | III | Colon | 50-59 | M | 2.6 | 0.196 | 17.05 ± 2.83 | 0.01 |
| C6 | III | Colon | 60-64 | F | 3.1 | 0.519 | 20.17 ± 0.71 | 0.01 |

| Sample ID | Cancer stage | Site of primary cancer | Age (yrs) | Sex | cfDNA stock conc. measured using Nanodrop (ng/µL) | cfDNA stock conc. measured using Qubit (ng/µL) | \|ΔZ\| at 100 kHz (ohms) | \|Δθ\| at 100 kHz (deg) |
|---|---|---|---|---|---|---|---|---|
| C7 | I | Rectum | 65-69 | M | 2.9 | 0.298 | 22.29 ± 0.00 | 0.00 |
| C9 | II | Colon | 60-64 | M | 2.2 | 0.141 | 15.15 ± 0.00 | 0.01 |
| C10 | II | Colon | 50-54 | F | 2.4 | 0.398 | 24.42 ± 0.00 | 0.02 |
| C11 | IV | Colon | 60-64 | M | 12 | 0.472 | 26.59 ± 0.71 | 0.03 |
| C12 | II | Colon | 65-69 | F | 2.1 | 0.24 | 17.16 | 0 |
| C13 | III | Colon | 50-54 | F | 2.4 | 0.672 | 18.79 | 0 |
| C14 | IV | Colon | 65-69 | M | 2.6 | 1.64 | 28.05 | 0.02 |
| C15 | III | Rectum | 65-69 | M | 2.3 | 0.568 | 37.72 | 0.02 |
| C16 | III | Colon | 50-54 | M | 2.9 | 1.44 | 16.57 | 0.01 |
| C17 | I | Colon | 65-69 | F | 2.7 | 0.688 | 17.91 | 0.01 |
| C18 | III | Rectum | 60-64 | M | 5.7 | 2.18 | 15.59 | 0.04 |
| C19 | II | Colon | 55-59 | F | 2.1 | 0.25 | 15.02 | 0 |
| C20 | I | Colon | 70-74 | F | 3.1 | 0.618 | 37.86 | 0.01 |
| C21 | IV | Colon | 50-54 | F | 2.8 | 1.48 | 10.37 | 0.01 |
| C22 | IV | Colon | 55-59 | M | 3.0 | 0.38 | 15.58 | 0.03 |
| C23 | II | Colon | 55-59 | F | 2.3 | 0.421 | 12.21 | 0.01 |
| C24 | IV | Colon | 75-79 | F | 3.2 | 1.18 | 13.5 | 0.01 |
| C25 | III | Colon | 60-64 | F | 3.3 | 1.36 | 13.7 | 0.03 |
| C26 | II | Colon | 76 | M | 1.3 | | 7.78 ± 1.07 | |
| C27 | IV | Colon | 57 | F | 3.6 | | 13.39 ± 0.18 | |
| C28 | III | Colon | 53 | M | 1.8 | | 1.72 ± 0.02 | |
| C29 | II | Colon | 72 | M | 5.7 | | 18.23 ± 0.54 | |
| C35 | IV | Colon | 55 | F | 36.4 | 21.5 | 12.02 | 0.05 |
| C36 | II | Rectum | 60 | F | 3.3 | 0.509 | 17.15 | 0.02 |
| C37 | I | Colon | 75 | F | 2.3 | 0.389 | 27.19 | 0.02 |
| C38 | IV | Colon | 53 | F | 3.9 | 0.943 | 3.89 | 0.01 |
| C39 | II | Colon | 65 | M | 2.2 | 0.718 | 11.97 | 0.02 |

| Sample ID | Cancer stage | Site of primary cancer | Age (yrs) | Sex | cfDNA stock conc. measured using Nanodrop (ng/µL) | cfDNA stock conc. measured using Qubit (ng/µL) | \|ΔZ\| at 100 kHz (ohms) | \|Δθ\| at 100 kHz (deg) |
|---|---|---|---|---|---|---|---|---|
| C40 | IV | Colon | 75 | M | 2.3 | 0.756 | 23.87 | 0.01 |
| C41 | IV | Colon | 70 | M | 2.5 | 0.626 | 9.14 | 0.01 |
| C42 | IV | Rectum | 65 | M | 2.7 | 0.719 | 17.54 | 0 |
| C43 | II | Colon | 71 | M | 4.4 | 1.55 | 22.22 | 0.04 |
| C44 | I | Rectum | 63 | F | 4.6 | 1.14 | 28.53 | 0.01 |
| C45 | III | Rectum | 57 | M | 6.9 | 2.19 | 15.84 | 0.02 |
| C46 | II | Colon | 47 | F | 4.5 | 1.47 | 12.6 | 0.02 |
| L1 | II | Colon | 54 | F | 4.9 | 0.955 | 23.2 | 0.03 |
| L2 | III | Rectum | 61 | M | 2.7 | 0.484 | 5.8 | 0.04 |
| L3 | II | Colon | 70 | M | 5.7 | 1.13 | 12.36 | 0.02 |
| L4 | I | Rectum | 62 | F | 2.2 | 0.611 | 15.15 | 0.01 |
| L5 | III | Rectum | 73 | M | 4.2 | 0.493 | 14.6 | 0.10 |
| L6 | II | Colon | 59 | M | 8.6 | 5.78 | 13.74 | 0.11 |

**Table S3.** Clinical information of healthy individual samples used for cfDNA extraction. Information where not available is left blank.

| Sample ID | Age (yrs) | Sex | cfDNA stock conc. measured using Nanodrop (ng/µL) | cfDNA stock conc. measured using Qubit (ng/µL) | \|ΔZ\| at 100 kHz (ohms) | \|Δθ\| at 100 kHz (deg) |
|---|---|---|---|---|---|---|
| H1 | 56 | M | 4.6 | 0.144 | 52.5 ± 2.83 | 0.01 |
| H2 | 63 | F | 3.3 | 0.238 | 45.17 ± 0.71 | 0.01 |
| H3 | 68 | F | 4.8 | 0.418 | 30.36 ± 0.71 | 0.02 |
| H4 | 59 | F | 3.8 | 0.336 | 31.62 ± 1.41 | 0.01 |
| H5 | 58 | M | 3.9 | 0.809 | 34.69 ± 0.71 | 0.02 |
| H6 | 54 | F | 3.2 | 0.287 | 32.69 ± 0.22 | 0.05 |
| H7 | 45 | M | 2.9 | 0.223 | 35.99 ± 0.71 | 0.01 |
| H8 | 60 | M | 3.1 | 0.368 | 38.27 ± 0.00 | 0.02 |
| H9 | 57 | M | 3.1 | 0.479 | 46.97 ± 5.66 | 0.01 |
| H11 | 71 | M | 3.6 | 0.362 | 49.85 ± 6.36 | 0.05 |

| | | | | | | |
|---|---|---|---|---|---|---|
| H12 | 55 | F | 2.6 | 0.421 | 24.82 ± 0.71 | 0.03 |
| H13 | 54 | F | 3.8 | 0.246 | 41.06 ± 1.41 | 0.08 |
| H14 | 62 | M | 3.4 | 0.481 | 37.47 ± 0.71 | 0.02 |
| H15 | 48 | F | 3.5 | 0.318 | 46.14 ± 2.83 | 0.03 |
| H16 | 60 | F | 1.5 | 0.247 | 29.15 ± 0.00 | 0.03 |
| H17 | 63 | M | 5.2 | 0.497 | 35.41 ± 2.12 | 0.03 |
| H18 | 48 | F | 4.0 | 0.217 | 47.7 ± 5.66 | 0.04 |
| H19 | 51 | F | 4.8 | 0.193 | 58.33 ± 1.41 | 0.04 |
| H20 | 50 | M | 2.9 | 0.387 | 40.53 ± 1.41 | 0.04 |
| H21 | 54 | F | 1.7 | 0.417 | 32.94 ± 1.41 | 0.01 |
| H22 | 57 | M | 2.4 | 0.435 | 43.14 ± 2.83 | 0.03 |
| H23 | 44 | F | 3.0 | 0.297 | 34.81 ± 0.71 | 0.02 |
| H24 | 35 | F | 3.9 | 0.319 | 48.71 ± 0.71 | 0.04 |
| H25 | 50 | F | 1.4 | | 42.97 ± 0.96 | |
| H26 | 47 | M | 1.2 | | 35.76 ± 1.39 | |
| H27 | 77 | M | 2.1 | | 34.44 ± 1.71 | |
| H28 | 55 | F | 3.1 | | 40.36 ± 0.60 | |
| H29 | 75 | F | 2.8 | | 34.51 ± 0.94 | |
| H30 | 62 | F | 4.3 | | 36.82 ± 0.94 | |
| H31 | 66 | F | 3.3 | | 33.53 ± 0.74 | |
| H32 | 68 | M | 2.3 | | 37.18 ± 1.29 | |
| H33 | 57 | M | 2.6 | | 41.08 ± 0.68 | |
| H34 | 53 | M | 5.0 | | 18.09 ± 0.74 | |

**Table S4.** Samples with history of cancer.

| Patient clinical status | No. of patients with history of cancer | Past cancer type | Asima Rev outcome |
|---|---|---|---|
| Healthy | 0 | N/A | - |
| CRC stage I | C20 | Leukemia | FN |
| CRC stage II | 0 | N/A | - |
| CRC stage III | C1 | Skin | TP |
| CRC stage IV | C3, C14, C38, C41 | Lung (C3),<br>CRC (C14, C38, C41) | TP |

**Table S5.** Demographic and clinical characteristics of CRC patients in TCGA.

| | **Adj. Normal (N=45)** | **Cancer (N=413)** |
|---|---|---|
| Age | | |
| Mean (sd) | 68.84 (12.23) | 64.56 (13.02) |
| Median | 70.00 | 66.00 |
| Min - Max | 40.00 - 89.00 | 31.00 - 89.00 |
| Gender | | |
| female | 21 | 190 |
| male | 24 | 220 |
| unknown | 0 | 3 |
| Pathologic stage | | |
| Adj. Normal | 45 | 0 |
| Stage I | 0 | 61 |
| Stage II | 0 | 151 |
| Stage III | 0 | 124 |
| Stage IV | 0 | 54 |
| Stage Unk. | 0 | 23 |

**Table S6**. Process of analysis of TEM images using ImageJ to calculate cfDNA aggregates in CRC surveillance.

| (1) Open TEM Image in Fiji (ImageJ). | (2) Subtract image background to make image more uniform given varying background fluorescence. | (3) Adjust image threshold to create a binary mask (all values set to either 0 or 1, black or white). |
|---|---|---|
| 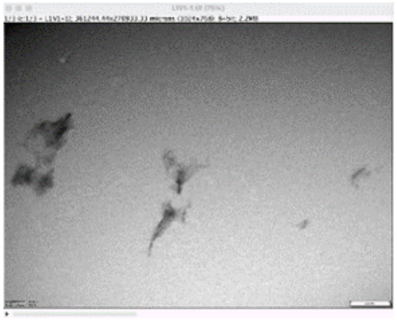 | 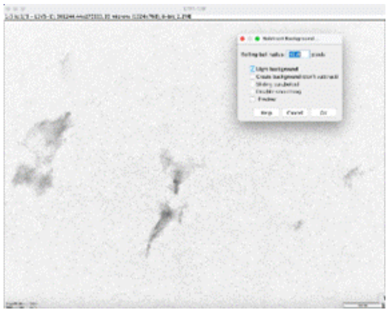 | 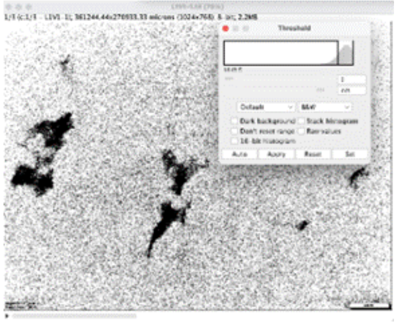 |

| (3) Erode then dilate the binary mask to remove the salt and pepper noise, then enhance the large pixel aggregates. | (4) Get an image histogram to derive the number of pixels at each binary value (0 and 1, black or white). | (5) Derive percentage of black pixels relative to all pixels, representing the percent area of the total image covered by aggregates. |
|---|---|---|
| 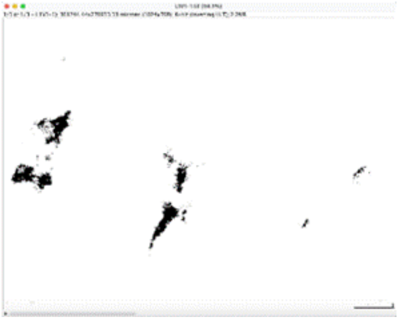 | 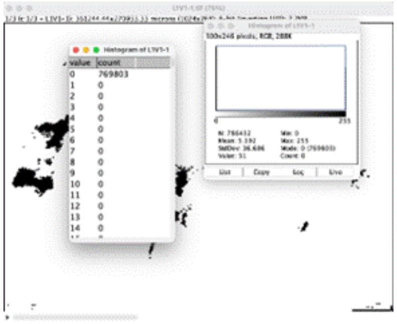 | No. white pixels: 769803<br>No. black pixels: 16629<br>No. total pixels: 786432<br>**Percentage black pixels or aggregates: 2.11** |

**Table S7.** Number of probes available and used for analysis in each dataset. 125111 OpenSea probes were common across all datasets.

| No. | Data Source | Total # of probes available | # of probes with 450k annotations | # of probes after filtering | # of probes annotated as OpenSea |
|---|---|---|---|---|---|
| 1 | TCGA-COAD and TCGA-READ | 486427 | 485512 | 398783 | 138091 |
| 2 | GSE131013 | 485577 | 485512 | 429944 | 152026 |
| 3 | GSE42752 | 485577 | 485512 | 429943 | 152026 |
| 4 | GSE48684 | 485577 | 485512 | 429944 | 152026 |
| 5 | GSE199057 | 865918 | 452453 | 400942 | 142532 |
| 6 | GSE240324 | 728310 | 369467 | 365981 | 130500 |
| 7 | GSE88824 | 485512 | 485512 | 429944 | 152026 |
| 8 | GSE42861 | 485577 | 485512 | 429944 | 152026 |
| 9 | GSE110185 | 866895 | 453093 | 401400 | 142664 |
| 10 | GSE186381 | 865859 | 452453 | 400962 | 142534 |
| 11 | GSE122126 | 423213 | 423213 | 398218 | 139885 |

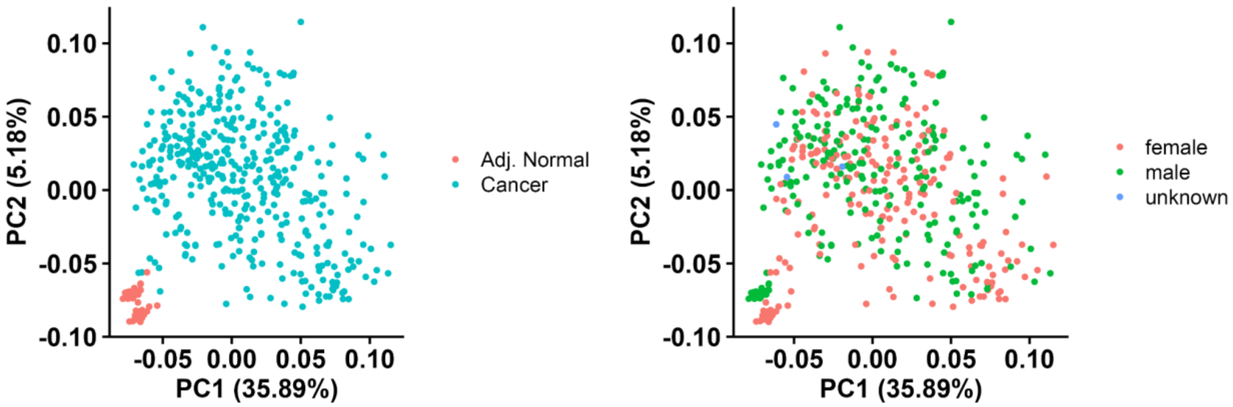


**Fig. S1.** Principal Components Analysis (PCA) plot of top 1000 most variable probes from 486, 427 total probes in TCGA without any filtering, colored by **(a)** cancer status and **(b)** gender.

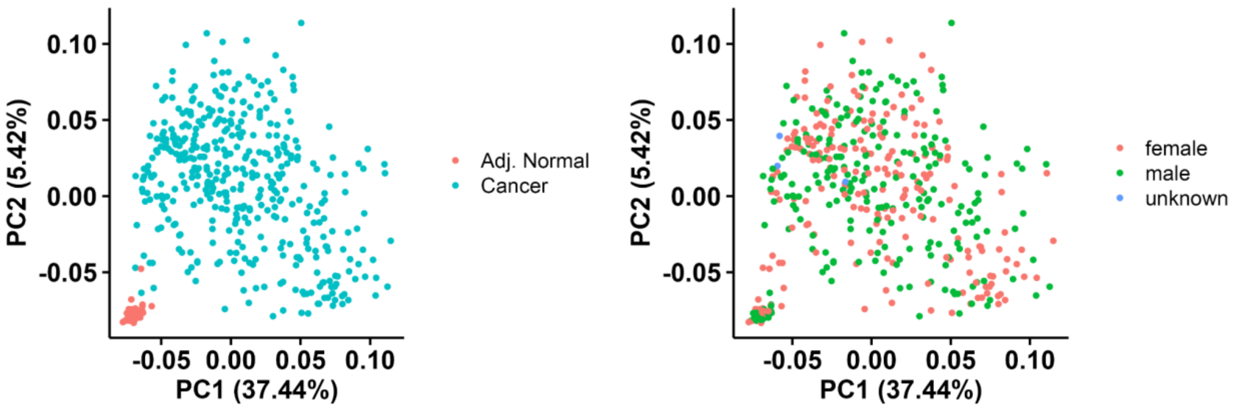


**Fig. S2.** PCA plot of top 1000 most variable probes from 398,783 total probes in TCGA after filtering, colored by **(a)** cancer status and **(b)** gender.

**Note S1. A discussion on why filtering of probes is required for Illumina Infinium methylation arrays.** Illumina Infinium methylation arrays use 50bp probes to query methylation status at various sites on the genome at a single base resolution. Before performing analysis on these data, it is common practice to filter out probes whose detection p-values are greater than a threshold, probes that occur in commonly known regions with single nucleotide polymorphisms (SNPs) that may affect CpG sites and probes that have shown to be cross-reactive. In addition, the PCA plots without removing probes on X and Y chromosomes showed that adjacent normal tissue samples cluster based on gender (Fig. S1), indicating that gender is a significant source of variation. Removing the probes from X and Y chromosomes led to cancer status being the main source of variance in the data (Fig. S2), enabling us to draw conclusions applicable to all samples.

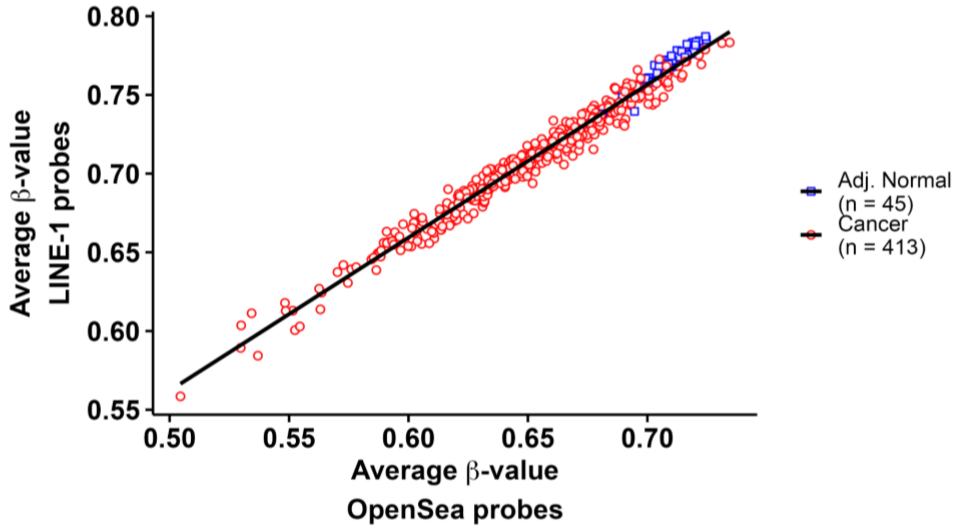


**Fig. S3.** Correlation between average β-value of LINE-1 probes and average β-value of OpenSea probes in TCGA.

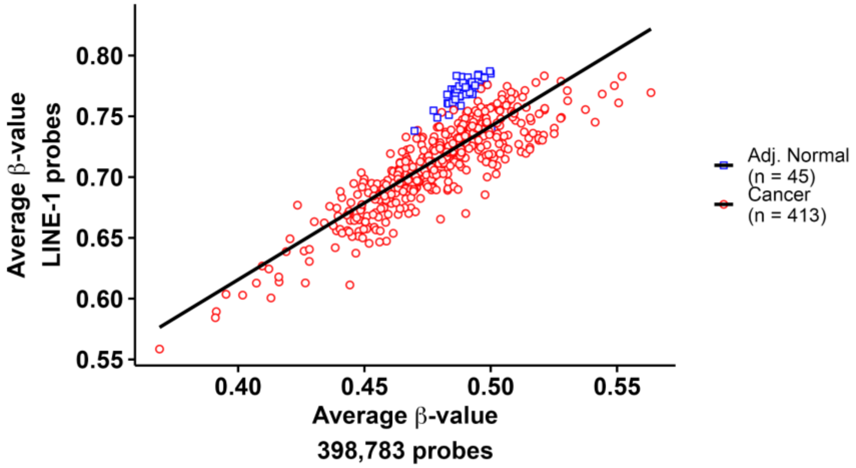


**Fig. S4.** Correlation between average β-value of all probes after filtering and average β-value of LINE-1 probes in TCGA.

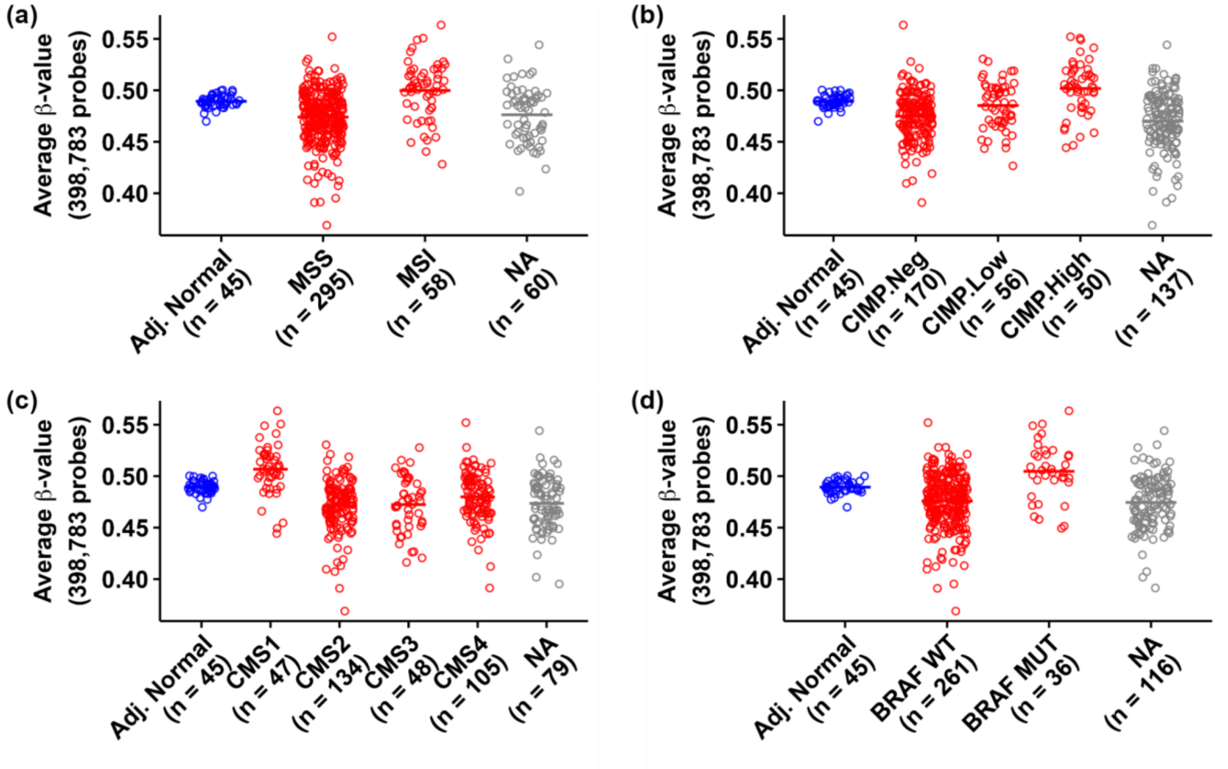


**Fig. S5.** Average β-value for all probes ($\beta_{av,\ all}$) in CRC patients with **(a)** microsatellite stability (MSS) and microsatellite instability (MSI); **(b)** CpG Island Methylator Phenotypes - High (CIMP.High), Low (CIMP.Low) and Negative (CIMP.Neg); **(c)** Consensus Molecular Subtypes (CMS) 1-4, and; **(d)** wild type (BRAF WT) and mutated (BRAF MUT) BRAF gene compared to adjacent normal tissue (Adj. Normal). NA refers to samples for which molecular subtype annotation was unavailable.

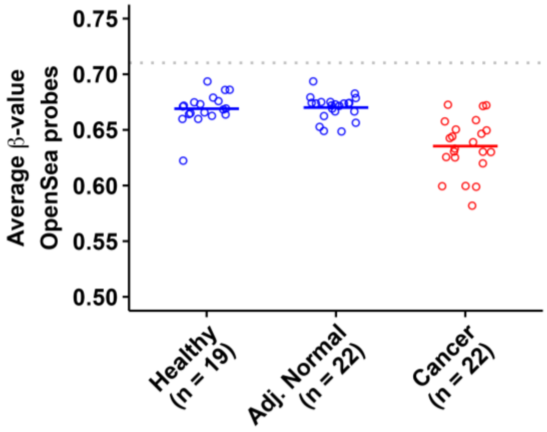


**Fig. S6.** Comparison of average β-values across OpenSea probes ($\beta_{av,\ OS}$) in healthy individuals (Healthy), adjacent normal tissue (Adj. Normal) from CRC patients, and CRC tissue (Cancer) in GSE42752. The gray line represents $\beta_{av,\ OS}$ in adjacent normal tissue from the TCGA study.

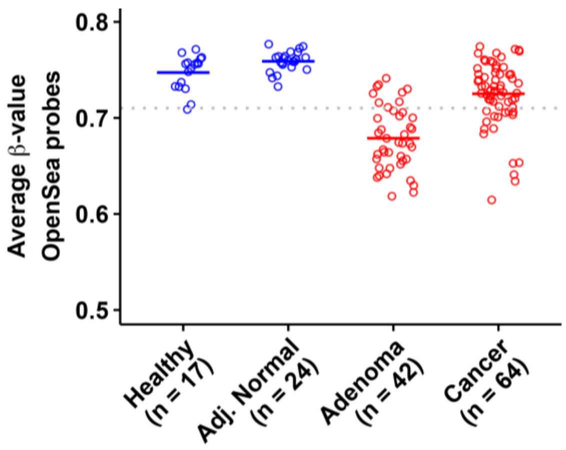


**Fig. S7.** Comparison of average β-values across OpenSea probes ($\beta_{av,\ OS}$) in healthy individuals (Healthy), adjacent normal tissue (Adj. Normal) from CRC patients, Adenoma patients (Adenoma) and CRC tissue (Cancer) in GSE48684. The gray line represents $\beta_{av,\ OS}$ in adjacent normal tissue from the TCGA study.

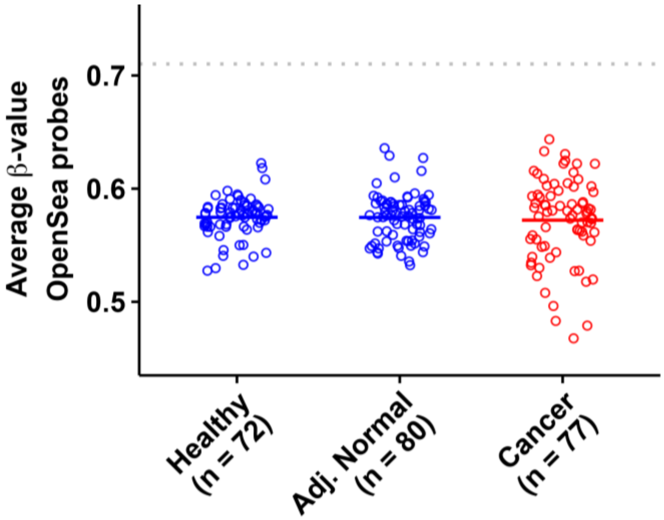


**Fig. S8.** Comparison of average β-values across OpenSea probes ($\beta_{av,\ OS}$) in healthy individuals (Healthy), adjacent normal tissue (Adj. Normal) from CRC patients, and CRC tissue (Cancer) in GSE199057. The gray line represents $\beta_{av,\ OS}$ in adjacent normal tissue from the TCGA study.

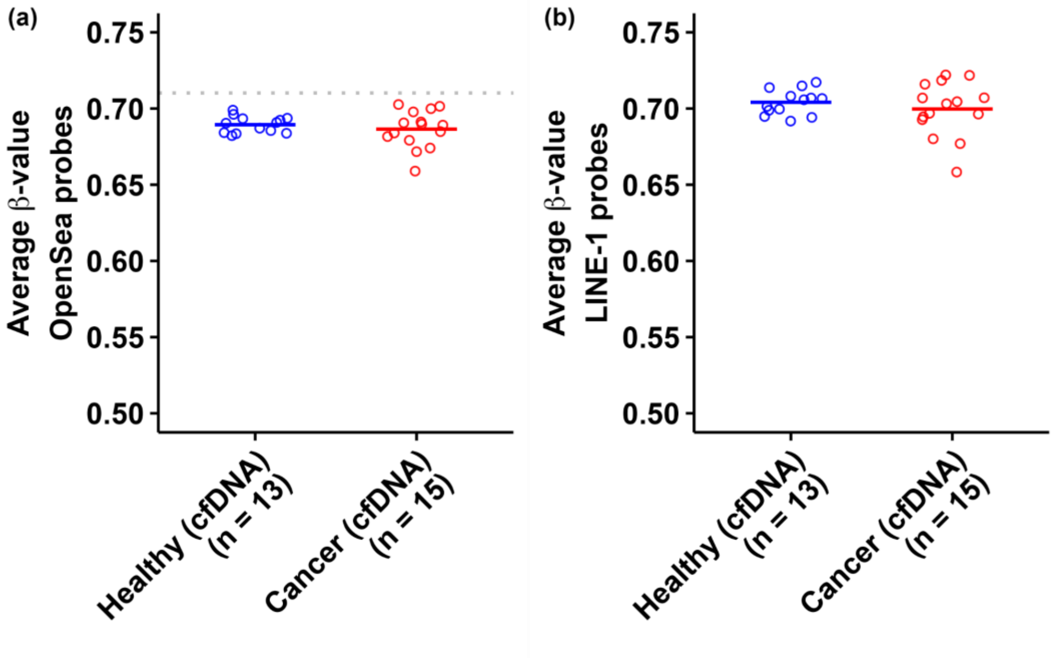


**Fig. S9.** Comparison of **(a)** average β-value across OpenSea probes ($\beta_{av,\ OS}$) and **(b)** average β-value across LINE-1 probes ($\beta_{av,\ LINE\text{-}1}$) in pooled cfDNA samples from healthy individuals and pooled cfDNA samples from CRC patients in GSE186381. The gray line represents $\beta_{av,\ OS}$ in adjacent normal tissue from the TCGA study.

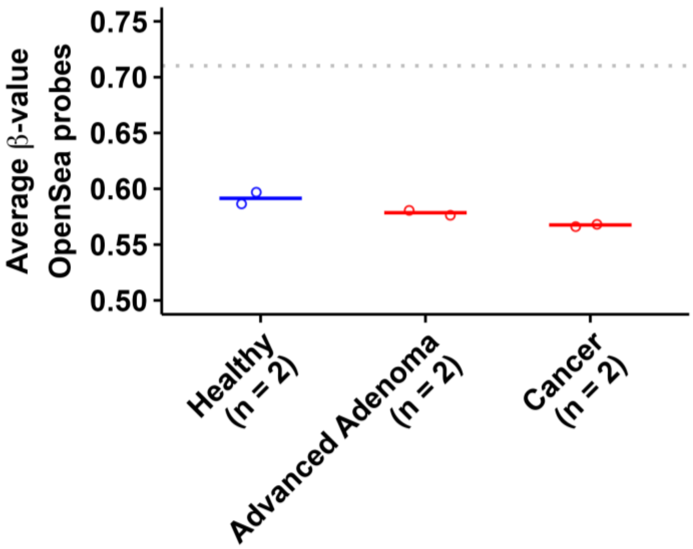


**Fig. S10.** Comparison of average β-value across OpenSea probes ($\beta_{av,\ OS}$) in pooled cfDNA samples from healthy individuals, pooled cfDNA samples from patients with advanced adenoma, and pooled cfDNA samples from CRC patients in GSE110185. The gray line represents $\beta_{av,\ OS}$ in adjacent normal tissue from the TCGA study.
